# The first photosynthetic mutant in diatoms targets the subunit γ of plastidial ATP synthase and reveals a key role of thylakoid electrochemical proton gradient in photosynthesis regulation and heterotrophic metabolism

**DOI:** 10.1101/2025.10.10.681070

**Authors:** Erik Jensen, Sien Audoor, Richard Kuras, Brignone Seleem, Alessandra Rogato, Giuliana D’Ippolito, Giovanna Benvenuto, Angelo Fontana, Yves Choquet, Benjamin Bailleul, Angela Falciatore

**Affiliations:** Laboratoire de Photobiologie et Physiologie des Plastes et des Microalgue, UMR7141, CNRS / Sorbonne Université, Institut de Biologie Physico-Chimique, Paris, France; Stazione Zoologica Anton Dohrn, Villa Comunale, Napoli, Italy; Institute of Bioscience and BioResources IBBR, National Research Council CNR, Via Pietro Castellino 111, 80131 Naples, Italy; Institute of Biomolecular Chemistry ICB, National Research Council CNR, Via Campi Flegrei 34, 80078 Pozzuoli, Italy; Department of Biology, University of Napoli “Federico II”, Via Cupa Nuova Cinthia 21, 80126 Napoli, Italy

**Keywords:** Diatom, Photosynthesis Mutant, Plastid ATP synthase subunit γ, proton motive force, photosynthetic control, phototroph and heterotroph metabolism, *Cyclotella cryptica*

## Abstract

Diatoms are major phytoplanktonic algae with secondary endosymbiotic plastids that differ in cellular and regulatory traits from those of the green lineage. Here we exploited the heterotrophic growth ability of *Cyclotella cryptica* to create the first diatom photosynthetic mutant by CRISPR-Cas inactivation of the nucleus-encoded ATP synthase subunit γ. These mutants showed impaired phototrophic capacity and altered thylakoids morphology. In absence of γ-ATP, protons that accumulate in the thylakoid lumen slow down the cytochrome *b_6_f* complex, thus keeping the electron carriers downhill oxidized. These results, reversible when the proton gradient is suppressed, demonstrate the existence of a photosynthetic control in diatoms. At variance with the wild type, γ ATP synthase mutants cannot grow heterotrophically in darkness nor in the light when photosystem II is inhibited. This requirement of heterotrophic growth on photosynthetic electron transfer or on the presence of plastidial ATP synthase suggests that the proton motive force (pmf) is a central integrator of the metabolic interaction between photosynthesis and heterotrophy. Our results establish *C. crytica* as a robust model for analysis of photosynthetic function, regulation and metabolic integration in organisms with secondary plastids.

**Teaser:** Mutagenesis in the facultative autotroph diatom *Cyclotella cryptica* enables exploration of essential plastid functions in diatoms

## INTRODUCTION

Diatoms are eukaryotic microalgae of the Stramenopile lineage thriving in all aquatic ecosystems and contributing to ∼20 % of Earth primary production (*1*). Derived from a series of secondary endosymbiosis events with both red and green algae (*2*, *3*), their plastids show distinct cellular features compared to those of green and red algae, glaucocystophytes and land plants (*4*) originated through primary endosymbiosis. Their plastids are surrounded by four membranes, instead of two, their thylakoids are organized in stacks of three layers, running parallel to the envelope membranes, and lack a clear grana and stroma lamellae distinction, although a spatial organization of photosynthetic complexes has been proposed (*5*). CO_2_ fixation (*6*, *7*) is enhanced in a pyrenoid enveloped by protein shells and possibly by the numerous physical contacts between plastid and mitochondria that could facilitate energy exchanges (*8*). While structures and functions of oxygenic photosynthesis complexes are conserved across phototrophs, diatoms also exhibit distinct acclimation strategies compared to green lineage models. Highly successful in turbulent environments (*9*), also thanks to very efficient yet simple photoprotective mechanisms (*10–12*) and light sensing strategies (*13*), diatoms also show a remarkable resilience to extreme environmental conditions, being able to survive under prolonged darkness during polar night or under sea-ice (*14*, *15*), or when buried under sediments.

Metabolic flexibility is likely another key factor in their ecological success. While most diatom species are obligate phototrophs, some are able of heterotrophic growth in the presence of different organic carbon sources (*16*, *17*), Additionally, certain species have been also described as mixotrophic, although typically only under specific conditions (*17*,*18*). The mechanisms controlling different trophic modes, as well as the impact of this regulation on diatom metabolism and growth are still largely unknown. This is in part due to the fact that, despite recent advances in diatom molecular biology, the mechanisms governing plastid biogenesis and *homeostasis*, as well as their cross-talk with other metabolic pathways and cell organelles remain largely unexplored. Moreover, many plastid-localized, nuclear-encoded factors, believed to regulate photosynthesis and plastid-related processes, still have unknown functions (*19*).

More than 80 years ago (*20–22*), the green alga *Chlamydomonas reinhardtii* emerged as a key model organism in plastid biology research, primarily due to its ability to grow both autotrophically and heterotrophically. This metabolic flexibility enabled researchers to generate and progressively study mutants deficient in photosynthesis and plastid biogenesis (*23–27*). To date, this strategy has not been feasible in diatoms, as the most established molecular models such as *Thalassiosira pseudonana* or *Phaeodactylum tricornutum*(*16*) are obligate phototrophs. A few studies have reported *P. tricornutum* mixotrophy growth with the supplement of glycerol, but only under nutrient-deficient conditions, which also negatively affect growth (*18*). Alternatively, a potential shift to heterotrophy has been observed in transgenic lines expressing an exogenous glucose transporter (*28*, *29*).

Here, we investigated the centric diatom *Cyclotella cryptica* as a promising model for studying diatom plastid biology. This species is one of the few laboratory-cultivated diatoms known to grow heterotrophically, using glucose as a source of reduced carbon (*30–32*). Moreover, it was the first diatom in which nuclear transformation was successfully achieved (*33*) and its genome has been recently sequenced and annotated (*34–36*), also due to its ability to accumulate triacylglycerols and other functional products under specific growth conditions and its potential in biotechnological applications (*37–41*).

Taking advantage of *C. cryptica* heterotrophic growth ability, we thus established genome editing by CRISPR-Cas9 in this species and generated the first photosynthesis diatom mutant targeting the nuclear-encoded *ATPC* gene, encoding the γ subunit of plastidial (CF1/Fo) ATP synthase. CF₁Fo ATP synthase is an enzyme that synthesizes ATP during photosynthesis by using the electrochemical proton gradient -or proton motive force (pmf)-generated across the thylakoid membrane during the light reactions of photosynthesis (*42*). It is made of two different sectors: the CFo part forms a membrane-embedded selective proton channel that allows protons to flow from the lumen to the stroma, while the CF₁ part uses this energy to catalyze the formation of ATP from ADP and inorganic phosphate. The flow of protons causes rotation of the central stalk, made of the subunits γ and ε within the catalytic head α_3_β_3_, driving conformational changes necessary for ATP synthesis (*42*, *43*). CFo is comprised of the AtpH, AtpI, AtpF and ATPG subunits in a 14:1:1:1 stoichiometry whereas CF₁ part is made of the subunits α, β, γ, 8 and ε in a 3:3:1:1:1 stoichiometry (reviewed in *44*, *45*). CF1 alone, or even a reconstituted α3β3y subcomplex, is able of ATP hydrolysis (without proton translocation), but ATP synthesis requires a fully assembled ATP synthase where the CF_1_ and CFo sectors are connected by the central stalk and the peripheral, made of ATPF and ATPG. This crucial γ subunit is, in diatoms, the only one encoded by the nucleus, at variance with the green lineage where ATPC (γ), ATPD (δ) and ATPG are nucleus-encoded.

The pmf plays a central role not only in ATP synthesis but also in various regulatory processes that enhance the robustness of photosynthesis under changing environmental conditions. Although its two components, Δψ (membrane potential) and ΔpH (proton gradient), are energertically equivalent in fueling ATP synthesis, they have distinct regulatory functions. In the green lineage (plants and green algae), ΔpH is a major regulator of photosynthesis. Thylakoid lumen acidification activates photoprotective mechanisms such as qE-type nonphotochemical quenching (NPQ) by protonating LhcSR3 or PsbS proteins and activating xanthophyll cycle de-epoxidase. This acidification also slows electron flow through cytochrome *b*_6_*f*, a process called “photosynthetic control”, thus protecting PSI by limiting over-reduction on the acceptor side, a condition that would otherwise leads to photodamage (*46–48*). This phenomenon is best visualized in ATP synthase mutants where proton accumulation in the lumen due to impaired proton efflux leads to an increased photosynthetic control under light (*46–48*). Beyond local regulation, ΔpH is linked to broader feedback mechanisms, including alternating electron flows (AEFs) or carbon concentrating mechanisms (CCMs) (*11*). Fine-tuning of pmf is therefore essential and occurs at two levels: adjustment between Δψ and ΔpH by ion channels, and a green lineage-specific redox control of ATP synthase regulating the overall amplitude. This redox control involves a conserved pair of cysteines in a 9-amino acid segment of the γ subunit, forming a disulfide bridge under the influence of light, which is reversibly reduced by thioredoxins (*46*, *49*, *50*). Finally, pmf also assists non-photosynthetic functions, such as the import of folded proteins into the thylakoid lumen via the Tat (twin arginine translocation) pathway, which requires a minimal pmf, regardless of photosynthetic efficiency. In line with this, ATP synthase is reversible, and can hydrolyze ATP (mainly supplied in the dark by mitochondria), pumping protons into the thylakoid lumen to maintain a minimal pmf in the dark (*50–52*).

The role of plastidial ATP synthase in regulating the pmf and the importance of this later in optimizing photosynthesis remain less well understood in diatoms than in the green lineage, although key differences are already apparent. In diatoms, the generation of pmf in the dark via the reversible activity of plastidial ATP synthase has been demonstrated, indicating a strong energetic coupling between the plastid and mitochondria (*8*). However, the precise control of pmf by CF₁F₀-ATPase differs markedly from that in green organisms, as diatoms lack the redox-sensitive cysteine pair responsible for ATP synthase regulation in the green lineage (*53*, *54*). Furthermore, the role of ΔpH as a master regulator is uncertain in diatoms (*11*); for example, their Lhcx proteins, homologs of PsbS/LhcSR3, do not respond to lumen acidification under high light, and their de-epoxidase enzyme function at near-neutral pH, unlike those of green algae and plants. Finally, it is still unclear whether photosynthetic control, as characterized in the green lineage, actually occurs in diatoms. Some diatoms of Nitzschia species have lost photosynthesis ability (*55*, *56*) and, consequently, also photosynthetic complexes, with the exception of ATP synthase (*56*, *57*). The retention of the plastidial ATP synthase in a heterotrophic diatom suggests that proton motive force across the thylakoid is required for other plastid metabolic activities related to heterotrophic metabolism. For all those reasons, ATP synthase mutants provide a valuable platform for uncovering regulatory roles of ATP synthase and pmf in diatom physiology and possible cross-talks between heterotrophic and phototrophic metabolisms. More generally, by establishing *C. cryptica* as a genetically viable model for the study of photosynthesis, this work paves the way for a better understanding of essential plastid functions in organisms with secondary endosymbiotic plastids.

## RESULTS

### Testing the potential of *C. cryptica* as a model for studying diatom photosynthesis

The ability of *C. cryptica* to undergo nuclear transformation and to grow heterotrophically made this species a promising candidate for the generation of photosynthesis mutants. Our first objective was to investigate the growth of *C. cryptica* under different trophic modes to identify heterotrophic conditions that could support the growth of photosynthetically deficient mutants (Fig. 1A). More specifically, we grew cells in liquid ESAW medium in continuous low or medium light (5 or 25 µmol photons m^-2^ s^-1^) or in complete darkness, with or without the addition of 10 mM glucose. We confirmed heterotrophy of *C. cryptica* as cells were able to grow in the dark in the presence of glucose, while, as expected, cells kept in the dark without glucose hardly performed one division (Fig. 1A). Moreover, the addition of a saturating concentration (1µM) of DCMU, an inhibitor of electron flow through photosystem II (PSII), completely abolished the growth of cells in darkness, low and medium light. Growth was however restored by supplementation with glucose. These effects were also observed at higher light intensity (50 µmol photons m^-2^ s^-1^, fig. S1). Together, these data demonstrate that glucose supplementation is enough to promote cell growth even in absence of PSII activity, a feature essential to inactivate genes essential for photosynthesis by mutagenesis.

**Fig. 1.**
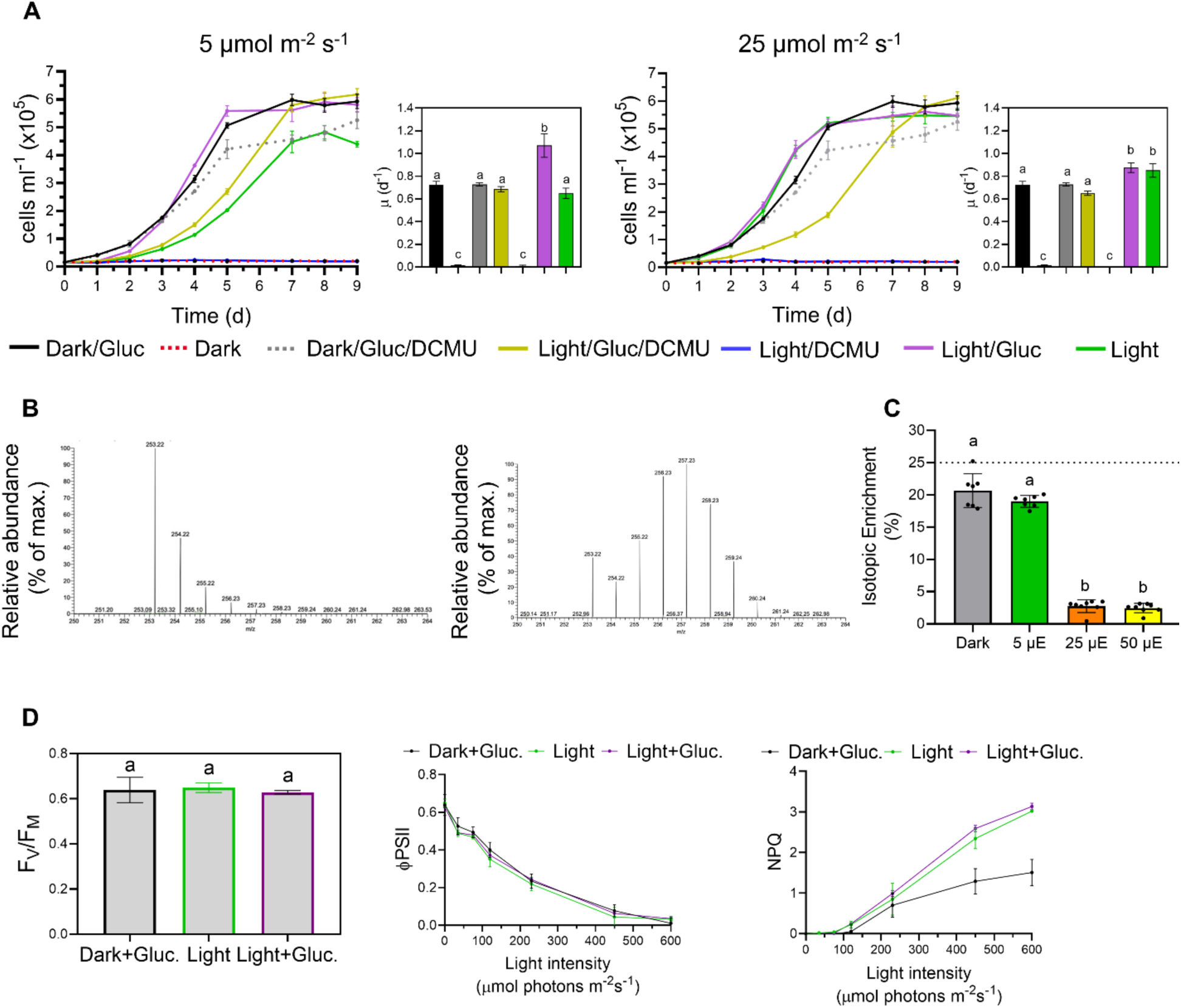
Characterization of the physiology of *C. cryptica* cultivated in different growth conditions. (**A**) *C. cryptica* wild type was grown in ESAW medium at light intensities of 5 µmol m^-2^ s^-1^ (left panel) or 25 µmol m^-2^ s^-1^ (right panel), with or without 10 mM glucose (gluc) and with or without the addition of 1 µM DCMU. Dark, dark + glucose and dark + glucose + DCMU controls are included in both panels. Cell growth was followed over time and growth rates (µ) were calculated from the slope of the linear logarithmic growth at the exponential phase (Fig. S1) and are shown as bar graphs. (**B**) MS spectrum of palmitoleic acid (C16:1 ω-7) from cells grown in the presence of glucose either at 25 µmol m^-2^ s^-1^ (left panel) or in the dark (right panel); (**C**) Heterotrophic contribution to central carbon metabolism was determined by percentage of ^13^C enrichment into palmitoleic acid from glycolytic metabolism of uniformly 2-^13^C-labelled glucose under different light regimes. The black dotted line indicates the maximum theoretical incorporation expected for pure heterotrophic growth. (**D**) Chlorophyll fluorescence analysis of cells grown in dark + glucose, light (25 µmol m^-2^ s^-1^) or light + glucose showing maximum quantum yield of PSII: (F_V_/F_M_, left panel), PSII operating efficiency: (<Ι_PSII_, middle panel) and non-photochemical quenching: N(PQ, right panel). All graphs are shown as the mean ± SD of three biological replicates. For bar graphs statistics, two-way ANOVA was used, with Tukey’s post-hoc analysis for multiple comparisons, where different letters represent significant differences (*p*≥0,05).

The addition of glucose did not affect the maximal growth rate of cells grown under medium light (Fig. 1A) or higher light (fig. S1B). This could be consistent with previous reports showing inhibition of glucose uptake in *C. cryptica* under moderate to high light (*31*). By contrast, under 5 µmol photons m^-2^ s^-1^, the addition of glucose boosted growth compared to purely phototrophic growth (Fig. 1A). The higher growth rate in the presence of light and glucose, compared to light or glucose alone, may indicate a concerted contribution of both photosynthesis and heterotrophic metabolism. To test this hypothesis we analyzed contribution of glucose-derived carbon to the *de novo* fatty acid biosynthesis via the plastidial fatty acid synthase (FAS) complex. To this end, *C. cryptica* cells were cultured in the presence of uniformly 2-^13^C-labeled glucose, which generates one molecule of 1-¹³C-labeled acetyl-CoA and one molecule of unlabeled acetyl-CoA per glucose molecule, irrespective of whether glucose is metabolized through the canonical Embden–Meyerhof–Parnas glycolytic pathway or through alternative routes such as the Entner–Doudoroff or pentose phosphate pathways. As shown for palmitoleic acid (C16:1) (Fig. 1B, left panel), mass isotopic distribution exhibited an isotopologue abundance consistent with a purely photosynthetic incorporation of environmental CO₂. In contrast, cells grown with 2-¹³C-glucose in complete darkness displayed isotopologue patterns indicative of extensive ¹³C incorporation derived from labeled glucose (Fig. 1B, right panel). The isotopologue distribution followed a Gaussian pattern, where progressive shifts toward higher *m/z* values reflected increasing ¹³C incorporation and, consequently, a greater contribution from heterotrophic metabolism.

Mass spectrometry (MS)-based analysis of isotopologue distributions across the major fatty acid species of *C. cryptica* revealed cumulative patterns consistent with these trends. Under irradiance levels ≥25 µmol photons m⁻² s⁻¹, supplementation of 1-^13^C glucose did not enhance significantly ¹³C labelling in fatty acids (fig. S2 and S3). In contrast, fatty acids from cells grown on this substrate at low light intensity (5 µmol photons m⁻² s⁻¹) or in complete darkness exhibited isotopic MS profiles characterized by a high relative abundance of labeled species (fig. S4 and S5). Given that only one of the four carbon atoms from each glucose molecule entering the central carbon metabolism carries a ¹³C label, the maximum theoretical enrichment of fatty acids is 25%. Analysis of relative isotopologue abundances (Fig. 1C) showed that ¹³C enrichment accounted for more than 23% of total fatty acid carbon in darkness and approximately 22% at 5 µmol photons m⁻² s⁻¹, both values approaching the theoretical maximum. In contrast, enrichment remained below 2% under the highest light conditions.

### The photosynthetic apparatus of *C. cryptica* is stably maintained in the dark and/or in the presence of glucose

This ability of *C. cryptica* to grow heterotrophically, as well as to undergo nuclear transformation makes it a promising candidate for generating photosynthesis mutants. However, it was also necessary to verify that the photosynthetic apparatus was not degraded when the alga was grown either in the dark or in the presence of an external source of carbon. Therefore we evaluated the photosynthetic efficiency of *C. cryptica* under phototrophic (light, with or without glucose) and heterotrophic (dark, with glucose) conditions (Fig. 1D). Chlorophyll *a* fluorescence measurements were used to determine the maximal PSII efficiency in the dark (F_V_/F_M_), the operational efficiency under light (<λ_PSII_), and the photoprotection capacity by non-photochemical quenching (NPQ). Overall, photosynthetic performance, as shown by F_V_/F_M_ and Φ_PSII_ measurements, remained similar regardless of the grown conditions tested. The NPQ capacity was significantly reduced but still present and with a similar light dependence, in cells grown in dark heterotrophic conditions, probably due to rearrangement of pigment composition or changes in the expression of LHCX proteins in the dark (*58*, *59*). These results suggest that, while the extent of photoprotection differs across trophic modes, the core photosynthetic machinery is preserved. This resilience underscores the potential of *C. cryptica* as a robust model to study photosynthetic function and regulation by using photosynthesis mutants.

### Generation of a photosynthetic mutant lacking the nuclear-encoded γ-subunit of the plastidial ATP synthase

The above results prompted us to generate photosynthesis mutants of *C. cryptica*. We selected as target for Crispr-Cas9 editing the only nucleus-encoded subunit of the plastidial ATP synthase, the gamma (γ) subunit encoded by the *ATPC* gene (CCRYP_015031-RA in version 2 of the genome; g19148.t1 in Nenasheva et al, (*36*)). Two CRISPR-Cas9 guide RNAs were designed to target the *ATPC* gene between the first exon and first intron (Fig. 2A, fig. S6A). The two gRNAs were *in vitro* assembled into RNPs and co-transformed with a vector containing the *NAT* resistance cassette using biolistic bombardment (see methods). Fifteen nourseothricin resistant colonies were obtained after transformation and selection of cells in the presence of 10 mM glucose and low light (5 µmol photons m^-2^ s^-1^). From those, three, named *atpc-*2, *atpc-* 8 and *atpc-*9, showed a size-modified amplicon (fig. S6B), resulting after gene editing from three different mutations in the *ATPC* gene (Fig. 2A, fig. S6C). Amplicons larger than the wild-type one were observed in clones *atpc*-2 and *atpc*-9. Sequence analysis of the PCR products showed that in both cases a fragment encompassing a *NAT* expression cassette from the selection vector, of 2516 and 1551 bp respectively (Fig. 2A and fig. S6B), was inserted close to the cutting site of guide RNA 1. The mutant strain *atpc*-8 had the amplicon of the predicted size for a 171 bp deletion spanning the cleavage sites of the two gRNAs (i.e., three base pairs after the PAM sequence), as confirmed by sequencing, and was further chosen for the complementation analysis (see below). All three mutations resulted in aberrant translation products (fig. S6D).

**Fig. 2.**
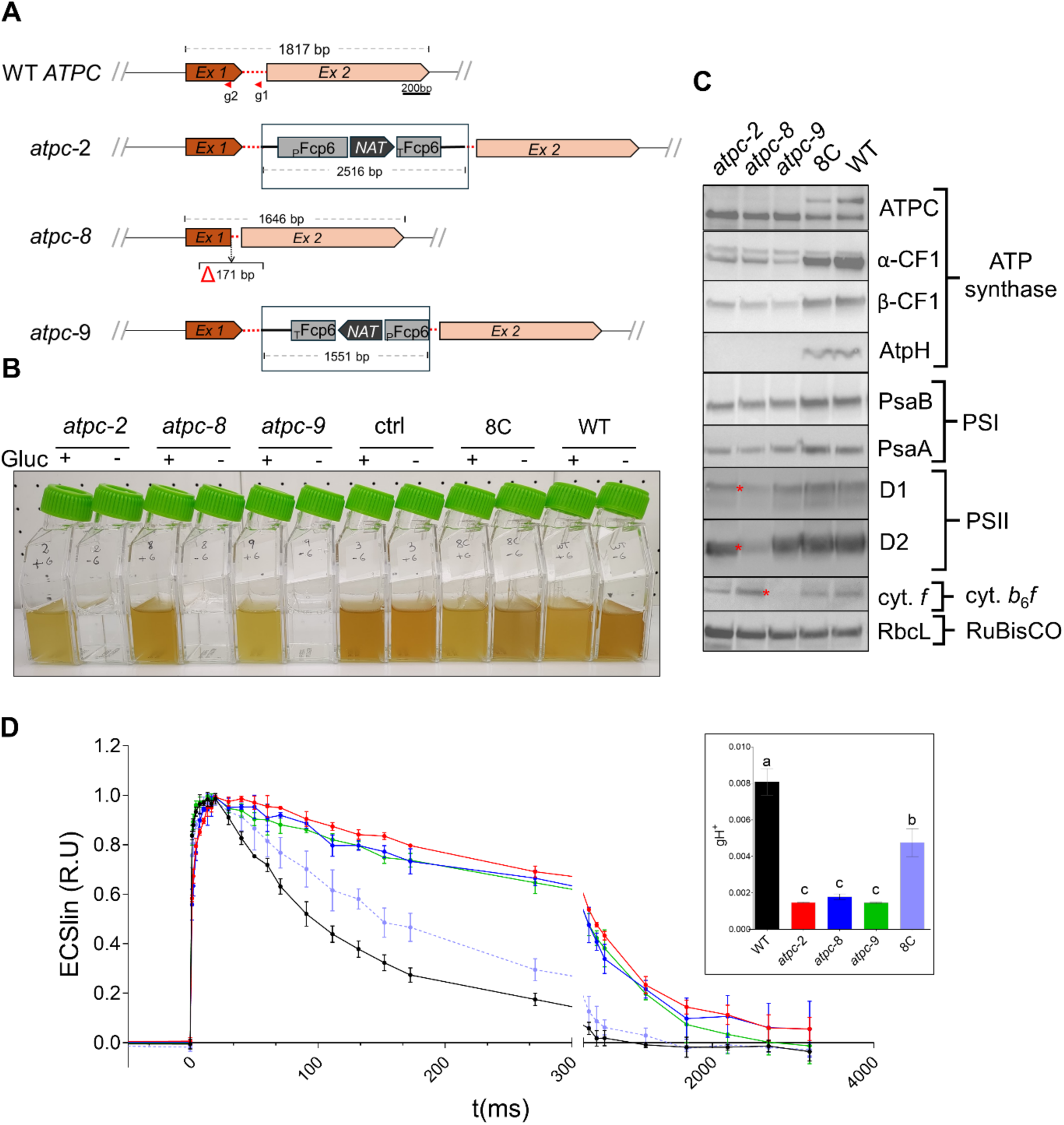
CRISPR-Cas9 mutagenesis of the *ATPC* gene and phenotype characterization of the resulting mutants. (**A**) Schematic representation *ATPC* locus (in scale) in the wild type (WT) and in the three independent mutant strains as the result of CRISPR-Cas9 editing. Exons 1 and 2 are shown as brown and pink arrows, respectively, and the intron as a red dotted line. The position of the two gRNA (g1 and g2) used for CRISPR-Cas9 gene editing is show with red arrowheads. Different size insertions of the NAT cassette in *atpc*-2 and *atpc*-9, coming from the co-transformed vector (see methods), are shown in boxes (not in scale). The deletion of 171 bp spanning the distance between the two gRNSs in *atpc*-8 is shown with a Δ symbol. In all three mutants, different truncated peptides are produced as result of a shift in the reading frame (for more details see text and Suppl. Fig. 6). (**B**) Growth the mutant strains in the presence or absence of glucose (10 mM) evidencing the loss of phototrophy. WT: wild-type strain, Ctrl: control strain that was transformed by proteolistic bombardment and selected on Nourseothricin-containing plates, but did not show a mutation in the *ATPC* gene, C8: Complemented *atpc*-8 strain. Independent flasks were inoculated at the same density (20 000 cells ml^-1^) and allowed to grow under 25 µmol m^-2^ s^-1^ with and without glucose (10 µM) for 7 days. (**C**) Immunoblot analysis of the accumulation of major photosynthetic subunits whole cell protein extracts using specific antibodies, as indicated on the right, in the wild-type, mutant and complemented strains. Red asterisks highlight proteins that are less expressed in one of the three mutants. **(D)** flash-induced ECS signal of mutants, wild-type and complemented 8C cells grown in low light (5 µmol m^-2^ s^-1^) in ESAW medium supplemented with 10 mM glucose. Boxed graph bar panel on the right shows the proton conductivity across the thylakoid membrane (gH^+^), extracted from the linear ECS exponential decay phase (see methods). Graphs show the average ± SD of three biological replicates. For statistics one-way ANOVA was used, with Tukey’s post-hoc analysis for multiple comparisons, where different letters represent significant differences (p≥0.05).

As expected for mutations preventing expression of the ATP synthase subunit γ, all mutant lines lost the ability to grow phototrophically. Therefore, they were kept growing under low light in the presence of 10 mM glucose (Fig. 2B). To further assess the implication of *atpc* mutation on the observed phenotypes, we complemented the *atpc* mutation by transforming the *atpc*-8 strain with a 2480 bp PCR product encompassing 691 bp upstream of the initiation codon, the fulllength *ATPC* coding sequence and 493 bp downstream of the stop codon. Transformed cells, selected for the recovery of photosynthetic growth on ESAW plates devoid of glucose, were then transferred to liquid medium (fig. S7A). As expected, the recovered 5 strains showed by PCR both the mutated and wild-type *ATPC* amplicons (fig. S7B). One of them, hereafter named 8C, was further analyzed along with the 3 independent mutants and the wild-type strain to assess the molecular and physiological effects of the *atpc* mutations.

### Characterization of the *atpc* mutants

Western blot analysis of total cell extract confirmed the absence of the γ subunit in the three mutants, and a recovered accumulation in the complemented 8C line, albeit to a lower level than in the wild-type strain (Fig. 2C). Interestingly, all mutants also lacked accumulation of the CFo AtpH subunit, illustrating a concerted accumulation of the subunits from the same photosynthetic protein complex (Fig. 2C). As already observed in a similar mutant of the green alga *C. reinhardtii* (*42*, *60*), the abundance of the α and β CF1 subunits was strongly reduced but not abolished. By contrast the accumulation of RbcL, PsaA and PsaB, respectively used as proxies of the accumulation of RuBisCO and PSI, was only marginally reduced. Unexpectedly, cytochrome *f* level was unaffected in the *atpc*-2 and *atpc*-8 mutants but strongly reduced in the *atpc*-9 strain, while the accumulation of the D1 and D2 core subunits of PSII was moderately decreased in the *atpc*-2 and *atpc*-9 mutants but strongly decreased in the *atpc*-8 mutants. Interestingly, 8C also recovered a wild-type level of the PSII subunits D1 and D2. These differences between the mutants are, however, not surprising considering that biolistic transformation often results in multiple insertions of the selection marker in independent loci.

To support these results with functional data, we measured the rate of proton efflux across the thylakoid membrane in the wild type, the three *atpc* mutants and the complemented strain 8C. This was assessed via the electrochromic shift (ECS) of photosynthetic pigments (*61*), which reflects variations in the trans-thylakoidal electric field generated by the charge movements due to photosynthetic electron transport and ATP synthase activity. We first analyzed the ECS signals in order to show the proper deconvolution of the linear and quadratic ECS signals (see Methods), a peculiarity of diatoms (*8*). Following a saturating laser flash (see Methods), the linear ECS signal exhibits a characteristic biphasic rise, corresponding to photosystem activity (phase *a*) and cytochrome b*_6_f* complex function (phase *b*), followed by a relaxation phase that reflects proton efflux to the stroma, catalyzed by ATP synthase (phase *c*) (*62*). As expected for a mutant lacking the CF1 γ subunit, *atpc* strains showed a significantly slower decay of the ECS signal, with an approximately fivefold reduction in the proton conductivity gH^+^ (Fig. 2D and inset) compared to the wild type. In contrast, the 8C strain restored proton efflux capacity to an intermediate level, consistent with the partial restoration of the γ protein subunit expression (Fig. 2C).

Next, we performed MS/MS-based quantitative proteomics to further characterize the proteomes of wild-type and *atpc* mutant strains cultivated in the presence of glucose under continuous low light conditions. Differential expression analyses from two independent experimental series confirmed the absence of the plastidial γ-subunit in all mutants. In these strains the peptide coverage was minimal (<2%) and restricted to regions upstream of the respective mutations (fig. S6D). Because these analyses, like immunoblots (Fig. 2C), revealed mutant-specific alterations, we focused on the core set of common proteins that exhibited consistent deregulation across all three *atpc* mutants, indicative of a conserved response to defective ATP synthase (Table I). A core set of 47 proteins were consistently downregulated compared to the wild-type, including plastid-encoded ATP synthase and PSII subunits, in agreement with the Western blot analysis (Fig. 2C). They were all, but two, either plastid-encoded or plastid-targeted (Table I), indicating substantial perturbation of processes occurring in this organelle. Interestingly, 17 pigment-binding antenna proteins were down-regulated in the mutants, an effect not observed in the green alga Chlamydomonas (*63*). In contrast, the 53 upregulated proteins were mostly all of unknown function and displayed broader subcellular distribution patterns (Table II), suggesting cellular compensatory mechanisms extending beyond plastid-specific responses. One of the few annotated proteins corresponds to the MLRQ subunit of the NADH-ubiquinone reductase complex 1, a key component of the mitochondrial respiratory chain. Interestingly, this protein has been identified as one of the most expressed proteins in the heterotroph diatom *Nitzschia putrida*, which has undergone a secondary loss of photosynthesis (*64*). We then analyzed by transmission electron microscopy (TEM) cells grown in low light conditions in the presence of glucose to visualize the effect of the ATP synthase deficiency at cellular level (Fig. 3). In the wild type as in the complemented strain, thylakoids span the entire plastid in homogeneous stacks of three, parallel to the envelope membrane. A central pyrenoid, where RuBisCO is concentrated, occupies the center of the plastid and is crossed by specialized thylakoids (Fig. 3). In the *atpc-8* mutant, while the pyrenoid is still present although with a more irregular shape, thylakoids appear as disorganized, and with locally dense, apparently super-stacked regions.

**Fig. 3.**
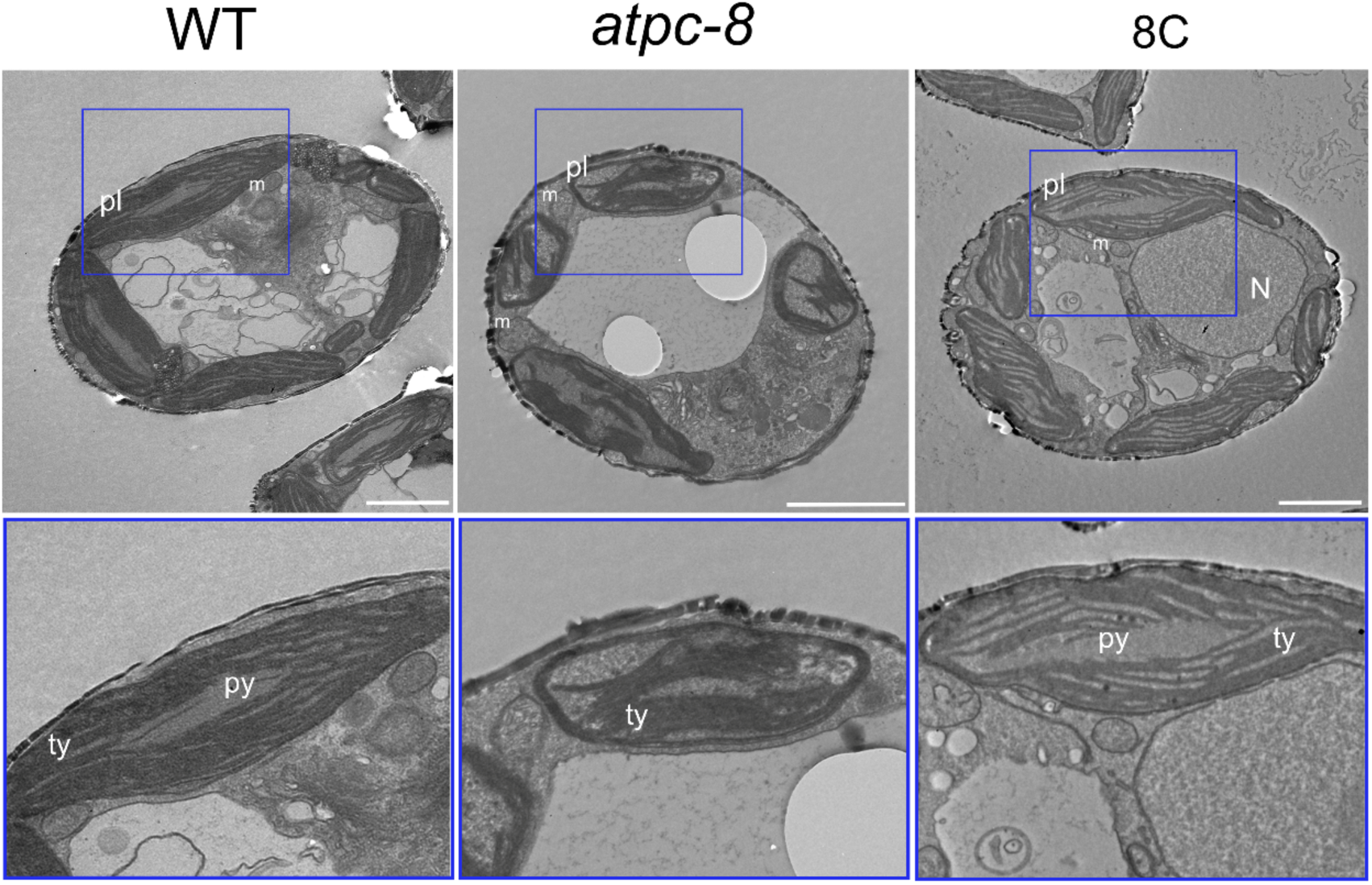
***atpc*-8 mutants have altered thylakoid architecture.** Transmission electron microscopy (TEM) of WT (left panels), *atpc-8* mutant (middle panels) and complemented line (8C; right panels). White bars correspond to 2 µm. Bottom panels are zoomed images of the blue squares shown in the panels above focusing on plastids. Pl: plastid, m: mitochondrion, N: nucleus, Py: pyrenoid, Ty: thylakoids.

**Table I.** List of differentially expressed down-regulated proteins in the three independent *atpc* mutants compared to the wild type.

| Down-regulated Protein |  |  |
| --- | --- | --- |
| name or ID* | Annotation | Gene location† |
| CpAtpE | AtpE | Pt |
| g20453.t1 | silicic acid transporter silicon transport | O |
| CpAtpG | AtpG | Pt |
| CpAtpF | AtpF | Pt |
| CpAtpD | AtpD | Pt |
| CpAtpA | AtpA | Pt |
| CpAtpB | AtpB | Pt |
| CpPsbD | PsbD | Pt |
| CpPsbA | PsbA | Pt |
| CpPsbC | PsbC | Pt |
| g16321.t1 | hypothetical protein | Pt |
| g19161.t1 | hypothetical protein | Pt |
| g19161.t2 | hypothetical protein | Pt |
| CpPsbE | PsbE | Pt |
| CpPsbB | PsbB | Pt |
| g17594.t1 | Rieske-like [2Fe-2S] domain | Pt |
| g14981.t1 | fucoxanthin chl ac light-harvesting protein major type; | Pt |
| CpPsbV | PsbV | Pt |
| g20683.t1 | fucoxanthin chlorophyll ac light-harvesting protein major type | Pt |
| g20683.t2 | fucoxanthin chlorophyll ac light-harvesting protein major type | Pt |
| g15561.t1 | fucoxanthin chlorophyll ac protein 2 | Pt |
| g15562.t1 | fucoxanthin chlorophyll ac protein 2 | Pt |
| g15567.t1 | fucoxanthin chlorophyll ac protein 2 | Pt |
| g19452.t1 | fucoxanthin chlorophyll ac protein 2 | Pt |
| g19454.t1 | fucoxanthin chlorophyll ac protein 2 | Pt |
| g19440.t1 | fucoxanthin chlorophyll ac protein 5 | Pt |
| g13955.t1 | fucoxanthin chlorophyll ac protein 8 | Pt |
| g13956.t1 | fucoxanthin chlorophyll ac protein 8 | Pt |
| g2143.t1 | fucoxanthin chlorophyll ac protein clade | Pt |
| g16996.t1 | fucoxanthin chlorophyll ac protein | Pt |
| g11872.t1 | fucoxanthin chl ac protein lhca clade | Pt |
| g20682.t1 | fucoxanthin chlorophyll ac light-harvesting protein major type | Pt |
| g20682.t2 | fucoxanthin chlorophyll ac light-harvesting protein major type | Pt |
| g26699.t1 | hypothetical protein; D-isomer specific 2-hydroxyacid dehydrogenase, NAD binding domain | Mt |
| g20684.t1 | fucoxanthin chlorophyll ac protein 6 | Pt |
| g20305.t1 | fucoxanthin chlorophyll ac light-harvesting protein | Pt |
| CpYcf46 | Ycf46 | Pt |
| CpPsaE | PsaE | Pt |
| CpPsaF | PsaF | Pt |
| CpPsaD | PsaD | Pt |
| CpPsaL | PsaL | Pt |
| g15753.t1 | hypothetical protein; Magnesium transporter NIPA | Pt |
| g17843.t1 | hypothetical protein; Bestrophin, RFP-TM, chloride channel | Pt |
| CpPsaA | PsaA | Pt |
| g7137.t1 | early light induced protein; Chlorophyll A-B binding protein | Pt |
| g13794.t1 | hypothetical protein | Pt |

\* Average coverage and score of proteins identified in the different mutants are indicated in Table S1.
†, Protein localization predicted by ASAFind: Pt, plastid, Mt, mitochondrion, O other localization.

**Table II.** List of up-regulated proteins in three independent *atpc* mutants compared to wild type.

| Up-regulated Protein |  |  |
| --- | --- | --- |
| name or ID* | Annotation | Gene location† |
| g3924.t1 | hypothetical protein | O |
| g13562.t1 | hypothetical protein | O |
| g7101.t1 | hypothetical protein | O |
| g7404.t1 | hypothetical protein | O |
| g9020.t1 | hypothetical protein | O |
| g15005.t1 | hypothetical protein | O |
| g2798.t1 | hypothetical protein | O |
| g25256.t1 | hypothetical protein; nitronate monooxygenase activity | SP |
| g13539.t1 | hypothetical protein | O |
| g17068.t1 | Metalloreductase STEAP4; cupric reductase activity | O |
| g14241.t1 | hypothetical protein; ShK toxin domain | SP |
| g2344.t1 | hypothetical protein | PPC |
| g23982.t1 | putative pyruvateferredoxin oxidoreductaseNADPH-cytochrome; sulfite reductase (ferredoxin) activity | O |
| g7537.t1 | hypothetical protein | O |
| g17724.t1 | Pyruvate formate lyase-like | PtL |
| g23152.t1 | hypothetical protein; Helicase associated domain | O |
| g14408.t1 | 4-hydroxyphenylpyruvate dioxygenase | O |
| g7994.t1 | hypothetical protein; Histone methylation protein DOT1 | O |
| g3305.t1 | hypothetical protein | PtH |
| g14851.t1 | hypothetical protein | O |
| g10495.t1 | Probable plastid-lipid-associated protein 8 chloroplastic; PAP fibrillin | PtH |
| g13973.t2 | hypothetical protein; Riboflavin kinase | O |
| g13973.t1 | hypothetical protein; Riboflavin kinase | O |
| g18821.t1 | hypothetical protein | O |
| g13482.t1 | hypothetical protein; AAA domain (dynein-related subfamily);g13482.t1\| hypothetical protein; AAA domain (dynein-related subfamily) | O |
| g69.t1 | hypothetical protein; prohibitin homologues | O |
| g26107.t1 | CUGBP Elav family member 6-like; RNA recognition motif. (a.k.a. RRM, RBD, or RNP domain) | O |
| g26962.t1 | hypothetical protein; AAA domain (dynein-related subfamily) | O |
| g26624.t1 | pyruvate phosphate dikinase; PEP-utilising enzyme, TIM barrel domain | O |
| g22806.t1 | hypothetical protein; Proline-rich nuclear receptor coactivator motif | O |
| g13482.t2 | hypothetical protein; AAA domain (dynein-related subfamily) | O |
| g16576.t1 | hypothetical protein | O |
| g12640.t1 | hypothetical protein | O |
| g7987.t1 | hypothetical protein | O |
| g5675.t1 | hypothetical protein; ABC transporter | O |
| g16346.t1 | hypothetical protein | O |
| g1891.t1 | hypothetical protein | O |
| g6905.t1 | hypothetical protein | O |
| g9260.t1 | hypothetical protein | O |
| g19303.t1 | hypothetical protein | O |
| g13030.t1 | hypothetical protein | O |
| g17854.t1 | hypothetical protein; Guanylate-binding protein, N-terminal domain | O |
| g6763.t1 | cyclin dependent kinase protein; Belongs to the protein kinase superfamily | O |
| g16319.t1 | hypothetical protein | PtH |
| g16319.t2 | hypothetical protein | PtH |
| g12584.t1 | hypothetical protein | PPC |
| g23475.t1 | hypothetical protein; KR domain | PtH |
| g6096.t1 | hypothetical protein; Serine aminopeptidase, S33 | Mt |
| g12494.t1 | hypothetical protein; NADH-ubiquinone reductase complex 1 MLRQ subunit | Mt |
| g15621.t1 | hypothetical protein | Mt |
| g18200.t1 | hypothetical protein | O |
| g12284.t1 | hypothetical protein; Prismane/CO dehydrogenase family | Mt |
| g6424.t1 | hypothetical protein | SP |

\* Average coverage and score of proteins identified in the different mutants are indicated in Table S1.
†, Protein localization predicted by ASAFind: Pt, plastid, Mt, mitochondrion, O other localization.

### Deletion of the γ-subunit impairs photosynthetic activity and reveals photosynthetic control in diatoms

We further investigated the effect of ATP synthase mutation on photosynthesis. The lack of proton efflux via ATP synthase in *atpc* mutants (Fig. 2 C) should result in proton accumulation in the thylakoid lumen under illumination. This was confirmed by measuring pH-dependent non-photochemical quenching (NPQ) in wild-type, mutant, and complemented strains under different light intensities (Fig. 4, left panel). Indeed, NPQ levels were significantly higher in mutants lacking the CF1 γ subunit compared to the wild-type and complemented 8C strains. As stated in the introduction, in plants and green algae, lumen acidification triggers photosynthetic control, slowing down the cytochrome *b*_6_*f* turnover (Fig. 5A). We used the *atpc-8* mutant to solve a long-lasting question (*10*) does the photosynthetic control operate in diatoms?

**Fig. 4.**
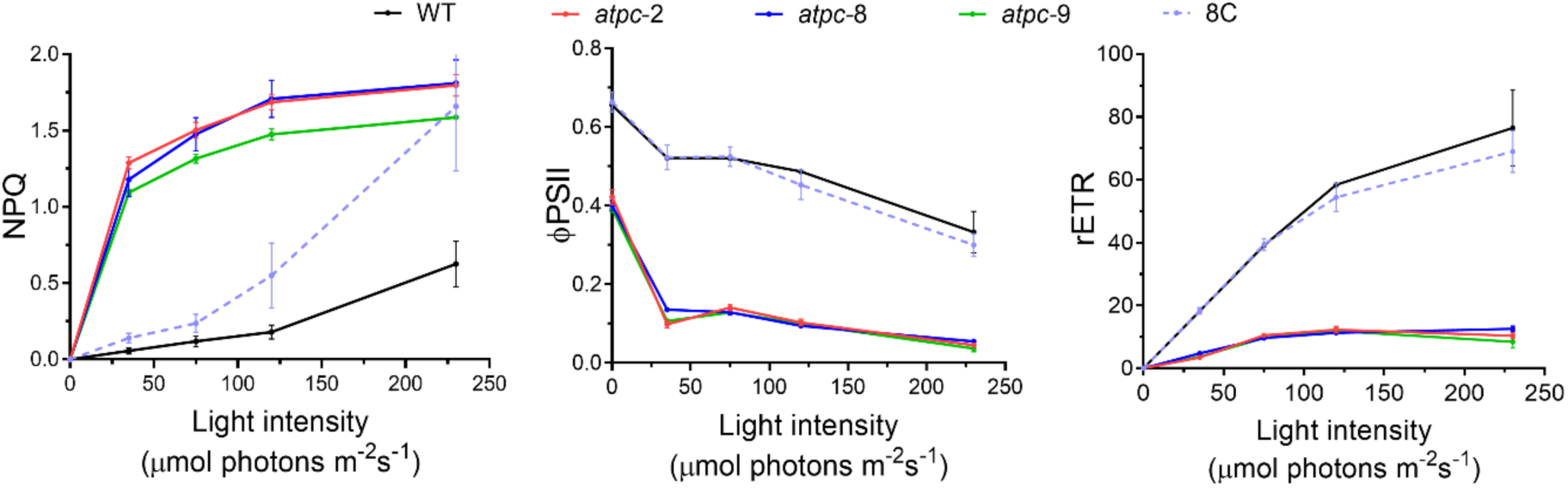
Effect of *atpc* mutation on photosynthetic efficiency. Non-photochemical quenching (NPQ; left panel), PSII operating efficiency (Φ_PSII_; middle panel), and relative electron transport rate (rETR; right panel) in mutants, wild-type (WT) and complemented 8C strain grown at 5 µmol photons m^-2^ s^-1^ in the presence of glucose are shown. All graphs show the average ± SD of three biological replicates.

**Fig. 5.**
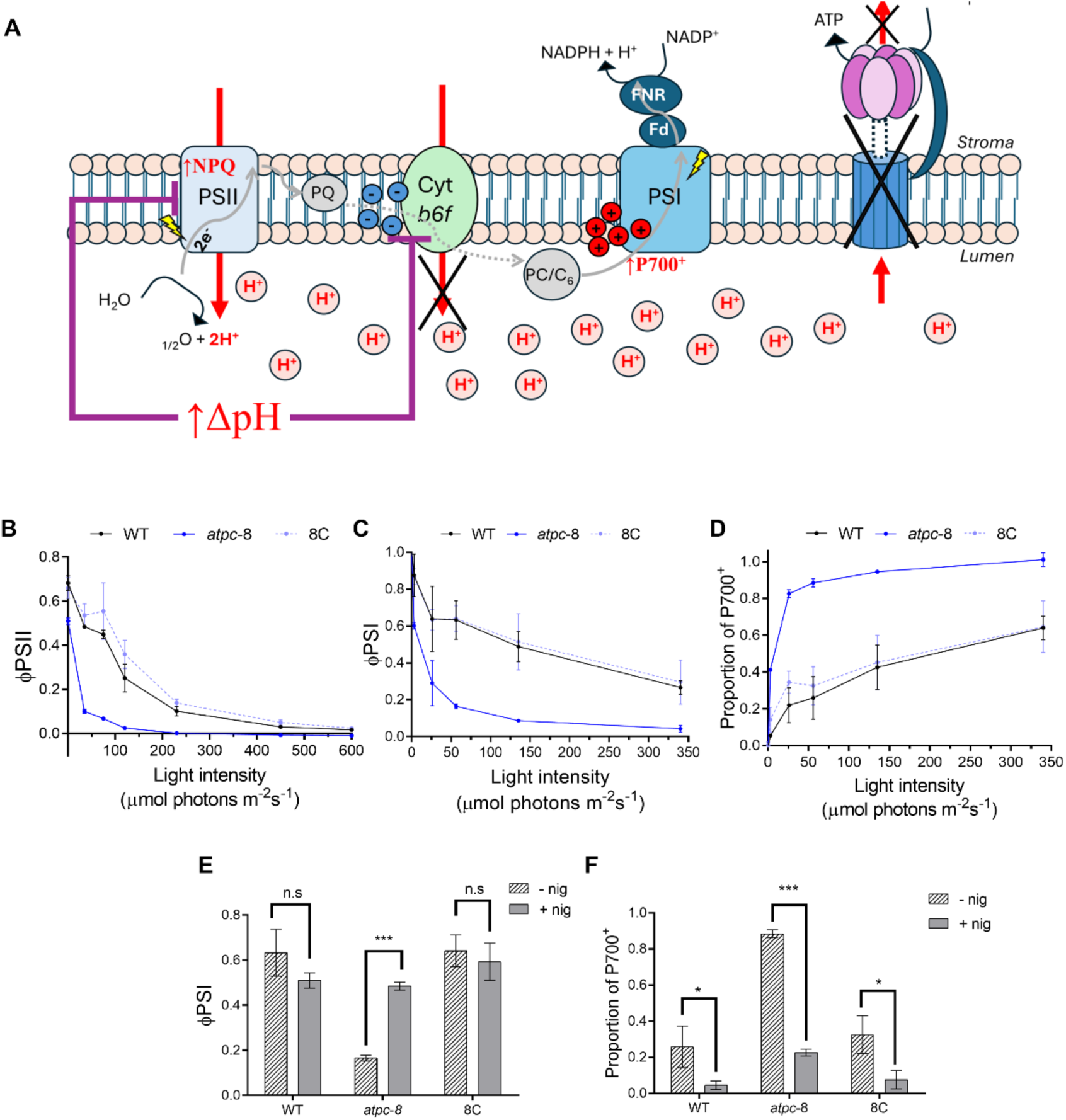
The *atpc* mutant allowed to confirm the existence of photosynthetic control in diatoms. (**A**) Scheme showing how photosynthetic control takes place in an ATP synthase-defective context. The inability to dissipate the proton gradient through ATP synthase leads to the acidification of the lumen, increasing the ΔpH across the thylakoid membrane. In consequence, electron flow slows down at the cytochrome *b_6_f* level, preventing over-reduction of downstream acceptors and photodamage of PSI. Lumen acidification is also associated with the induction of non-photochemical quenching (NPQ). Linear electron flow is shown as a grey arrow; with dotted line representing the slowed down electron flow. Proton flows are shown with red arrows. More detail in the main text. (**B**) PSII operating efficiency – <Ι_PSII_ – derived from chlorophyll *a* fluorescence analysis. (**C**) PSI operating efficiency – <Ι_PSI_) – extracted from P_700_ redox kinetics. (**D**) proportion of P_700_^+^, are shown in the wild-type (WT), *atpc*-8 and complemented 8C strains. (**E**) ΦPSI and (**F**) proportion of P_700_^+^ in cells exposed to 56 µmol m^-2^ s^-1^ in the presence of 1 µM nigericin. All graphs show the average ± SD of three biological replicates. t-test statistical analysis with pair-wise comparison was done in (E) and (F) (\**p*≥0.05, *** *p*≥0.001).

Suggesting the actual presence of photosynthetic control, photosynthetic electron transfer was significantly reduced in the mutant compared to the wild-type or complemented strains, at all light irradiances, as illustrated by the lower Φ_PSII_ and rETR values (Fig. 4, middle and right panels). To further confirm the occurrence of this regulation, we examined the redox state of electron transporters between cytochrome *b_6_f* Qo site and PSI, namely cytochrome *f* in cytochrome *b_6_f* complex, the electron shuttle c_6_ and P_700_, the PSI special pair. If photosynthetic control is induced in ATP synthase mutants, limiting electron transfer at the *b_6_f* complex, these transporters should remain significantly more oxidized in the mutants than in the WT. Indeed, in the *atpc*-8 mutant, the quantum yield of PSI (ΦPSI) mirrored the decline in Φ_PSII_, showing a dramatic reduction even at low light (Fig. 5B, C), confirming an impaired electron transport. Upon illumination (∼56 µmol photons m^-^² s^-^¹), we observed a higher oxidation level for P_700_ (Fig. 5D) and c-type cytochromes (fig S8) in the mutant, as expected from a high photosynthetic control in the mutant.

We reasoned that the dissipation of ΔpH should attenuate this control, restore electron flow and reduce P_700_ and c-type cytochromes. We thus treated cells with 1 µM nigericin, an H⁺/K⁺ antiporter that dissipates the proton gradient without affecting the Λλλ′ component of the pmf. Suppression of NPQ in the *atpc*-8 mutant confirmed the successful dissipation of ΔpH (fig. S96). After nigericin treatment, Φ_PSI_ in the mutant increased significantly in the *atpc*-8 mutant, approaching WT levels (Fig. 5E), while Φ_PSII_ also showed partial recovery (fig. S9). In contrast, nigericin had minimal effect on Φ_PSI_ and Φ_PSII_ in WT and 8C strains. Furthermore, nigericin treatment decreased the proportion of oxidized P_700_ (Fig. 5F). The observed reduction in electron flow (as shown by both Φ_PSI_ and Φ_PSII_) and the strong oxidation of the electron transport chain after cytochrome b*_6_f*, together with the fact that both of these effects are reversed when ΔpH is dissipated, provide definitive evidence that photosynthetic control is active in diatoms, just as it is in organisms of the green lineage.

### Heterotrophic growth would require the generation of PMF through photosynthetic electron transfer or the reverse activity of ATP synthase

To further explore the interaction between phototrophic and heterotrophic metabolisms, we assessed the impact of photosynthesis inhibition on growth by treating the different strains with DCMU, a PSII inhibitor, under various trophic conditions. As expected, in the wild type, DCMU had no effect on growth in total darkness and completely inhibited growth under purely phototrophic conditions (light without glucose; Fig. 1A), confirming the essential role of photosynthetic electron transfer under these conditions. At 50 µmol photons m⁻² s⁻¹ in the presence of glucose, DCMU treatment induced a slowed down but sustained growth, indicating that cells can switch to a heterotrophic metabolism when photosynthesis is blocked.

Interestingly, the response of the *atpc* mutants was very different: the addition of DCMU under mixotrophic conditions largely inhibited growth (Fig. 6A). These results reveal a strong dependence of heterotrophic metabolism on the photochemical phase of photosynthesis: it seems to require either the presence of the ATP synthase or photosynthetic electron transport. The nature of this control appears to be linked to the pmf. Indeed, pmf is the only photosynthesis-related parameter that can be generated either by photosynthetic electron transport in the light or by ATP hydrolysis by ATP synthase in the dark, using mitochondrial ATP (*8*, *10*). Such a model would predict that *atpc* mutants are unable to grow in the dark, even in the presence of glucose. To test this, we cultured *atpc* mutants in the dark, with or without glucose supplementation. As expected, in the absence of glucose, the mutants, like the WT mutants, divided only once before complete growth arrest (Fig. 6B). In the presence of glucose, the mutants could complete three divisions before their growth was fully arrested (Fig. 6B), unlike the WT and the complemented 8C line which displayed robust and sustained heterotrophic growth. This suggests that pmf is a key integrator of the metabolic dialogue, necessary for the activation or maintenance of heterotrophic metabolism.

**Fig. 6.**
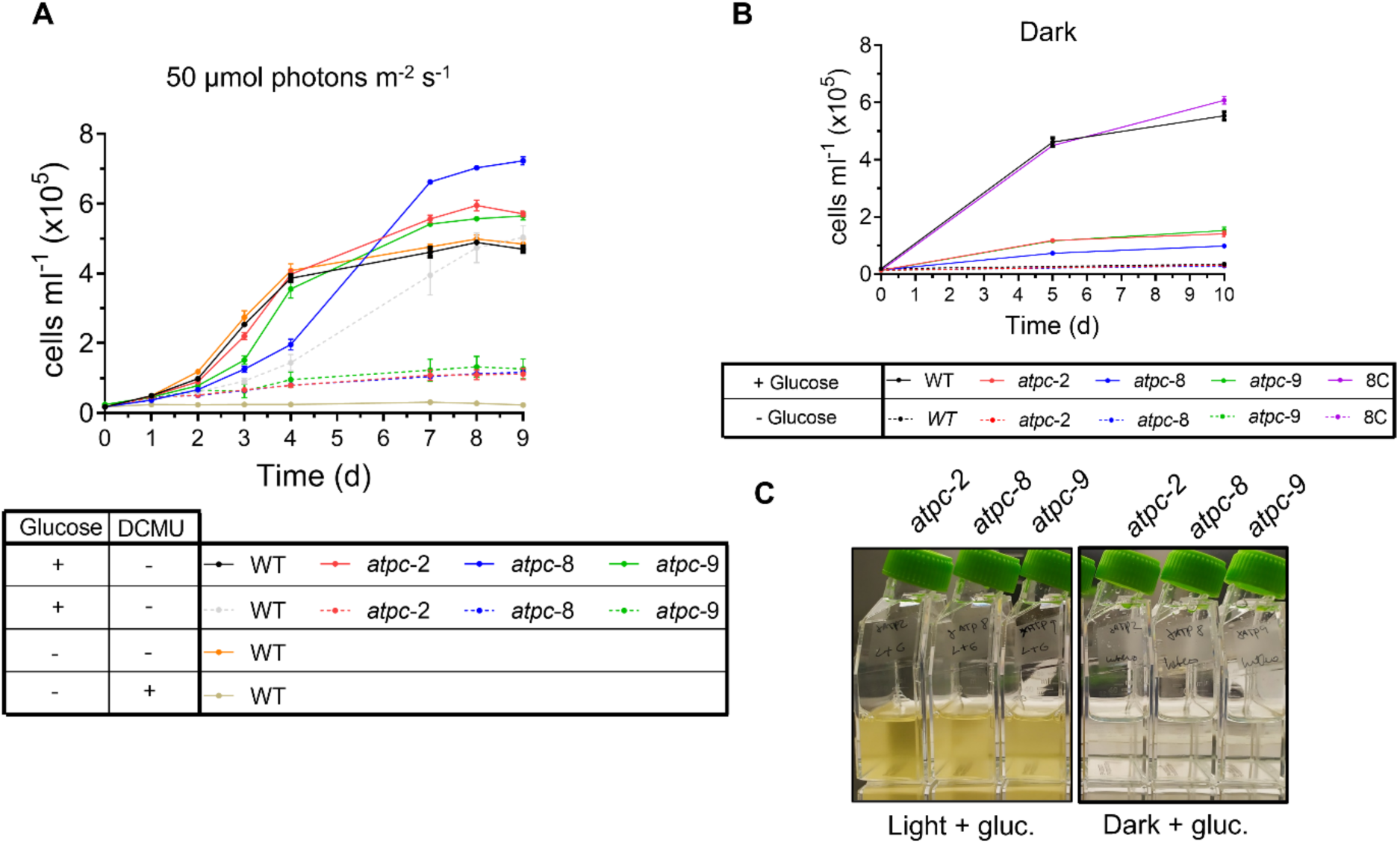
Effect of DCMU and of dark cultivation on mutant cell growth. (**A**) Growth curves of the three *atpc* mutants and wild-type under 50 µmol photons m^-2^ s^-1^. The addition of 10 mM glucose or 1 µM DCMU is indicated in the lower table. (**B**) The three *atpc* mutants, wild-type (WT) and Complemented strain (8C) were cultured with or without glucose in the dark for 10 days. (**C**) Representative pictures of the *atpc* mutants cultivated in medium supplemented with glucose under light or dark conditions. Growth curves are shown as the mean ± SD of three biological replicates.

## DISCUSSION

Over the past 20 years, advances in genomics and molecular biology have led to the establishment of new diatom experimental model systems, including *P. tricornutum* (*65*), *T. pseudonana* (recently renamed *Cyclotella nana*) (*66*), *Fragilariopsis cylindrus* (*67*), and *Pseudo-nitzschia multistriata* (*68*). Functional investigations in these algae are progressively uncovering cellular and metabolic processes and regulatory mechanisms that distinguish diatoms from green algae, plants, and animals (*69*). To uncover essential yet unknown processes related to photosynthetic activity, processes that cannot be studied in the other diatom models due to their strictly photoautotrophic metabolism, we established genome editing in the facultative heterotroph *C. cryptica*. By targeting the nucleus-encoded γ subunit of the plastidial ATP synthase by CRISPR/Cas-mediated genome editing, this study advances our understanding of diatom plastid biology.

The decreased accumulation of all plastid-encoded ATP synthase subunits observed in the ATP γ KO indicates a coordination between the nuclear and plastid genomes, as has already demonstrated in organisms derived from primary endosymbiosis (*42*, *70*, *71*). In the green lineage two major mechanisms account for the concerted accumulation of the different subunits of a same protein complex (*71*). Some subunits, particularly those encoded in the nucleus, are expressed normally but rapidly degraded when they cannot assemble, while many chloroplast-encoded subunits of the photosynthetic apparatus show an assembly-dependent regulation of their synthesis, a process known as the control by epistasy of synthesis (CES) process. In the absence of their assembly partners, the rate of synthesis of CES subunits is dramatically reduced. Described in the chloroplast of green algae and higher plants (*71*, *72*) as well as in the mitochondria of yeast(*71*), it remains to be determined whether similar CES process also exists in the plastids of diatoms. It cannot, however, account for the down regulation of PSII and antenna proteins, which seems to be diatom-specific, as ATP synthase mutants of Chlamydomonas do not show any alteration in the PSII and LHC content (*63*). This observation possibly suggests a retrograde signaling that will deserve further investigations.

Our analyses also showed a disorganization of the thylakoid membranes in the *atpc* mutants, with dense patches of apparently hyper-stacked thylakoids, while in the wild type stcks of 3 layered thylakoids run parallel to the envelope membranes. This is in contrast with the equivalent mutants of *C. reinhardtii* that, as long they are kept in low light, retain a normal chloroplast ultrastructure. Immunolocalization experiments in *P. tricornutum* showed that PSI and ATP synthase are mostly found in the external, stroma-facing thylakoids, while PSII is mostly embedded in the internal thylakoids (*5*, *73*). It is thus possible that the bulky ATP synthase complex protruding in the stroma and therefore segregated to the external thylakoids would be required to maintain the three-thylakoids organization.

The characterization of the γ ATP subunit also provided novel information on diatom photosynthesis regulation. As expected for a mutant lacking ATP synthase, our *atpc* mutants in *C. cryptica* show impaired photosynthesis. Moreover, they also showed decreased proton efflux to the stroma, the resulting proton accumulation leading to an increased non-photochemical quenching (NPQ). The over-accumulation of protons in the lumen observed in this mutant allowed us to test the existence of photosynthetic control in diatoms. The slowdown in electron transfer and oxidation of transporters between the cytochrome b*_6_f* complex and PSI, attenuated in the presence of an uncoupler, unambiguously demonstrates -for the first time-the existence of such a regulatory mechanism in diatoms. While this, at first glance, seems a mere confirmation of what is well established in the green lineage, it was legitimate to ask whether, in diatoms, the lumen could actually reach a pH low enough to trigger a slowdown of the b*_6_f* steps involving proton release. Indeed, diatoms exhibit unique characteristics when faced with lumen acidification: absence of Lhcx1 protonation, activation of de-epoxidase at near-neutral pH, etc. Too often, the “gray areas” of diatom physiology are filled by analogies with the green lineage, out of habit or due to a lack of specific data. Therefore, our results provide a clear answer to this question, and highlight the need to characterize the mechanisms specific to diatoms without systematically relying on paradigms from the green lineage.

Analysis of glucose metabolism under different light regimes demonstrates that *C. cryptica* possesses a pronounced capacity to modulate carbon metabolism in response to light availability, shifting from autotrophic CO₂ fixation to glucose-based heterotrophy under limiting irradiance (*31*, *32*, *74*). The marked ¹³C enrichment of fatty acids in low light and darkness confirms that glucose-derived carbon efficiently enters central carbon metabolism. Conversely, under moderate to high light intensities, glucose assimilation was almost completely suppressed, in agreement with early evidence of light-dependent inhibition of glucose uptake in this species (*31*). These findings showed that a transition from predominantly autotrophic to predominantly heterotrophic metabolism occurs between 25 and 5 µmol photons m⁻² s⁻¹. Above this threshold, cells remain photosynthetically active, with negligible glucose assimilation, even in the presence of glucose. Below 5 µmol photons m⁻² s⁻¹, metabolism is clearly dominated by glucose assimilation, though residual photosynthetic activity persists. This shift is in agreement with the observed effects on growth under combined low light and glucose availability.

Surprisingly enough, *atpc* mutants showed a conditional heterotrophic growth capacity. Able to successfully grow on external carbon source in the light, this ability is lost in the mutants in the presence of photosynthesis inhibitors, or when grown in the dark. On the contrary, the wild-type with a functional ATP synthase can grow on glucose, as soon as photosynthesis becomes limiting (i.e. under darkness or low light or in the presence of inhibitors). These results strongly suggest that the trans-thylakoidal pmf plays a role in the heterotrophic growth of *C. cryptica* cells. We found that under all conditions where a significant pmf was generated, either by photosynthesis (e.g., in wild-type or mutant cells exposed to light) or by reverse ATP synthase activity in the dark (in WT cells growing on glucose), cells were able to grow. In contrast, when pmf generation was impaired; i.e. in mutants kept in the dark or in the presence of DCMU under illumination, cells failed to grow. This highlights the requirement of heterotrophic metabolism for a pmf across the thylakoid. Many vital functions occurring in the plastid require energy, it is reasonable to believe that some of those vital functions require the energization of the thylakoid membrane; however, we have not yet identified the mechanism(s) at play.

Recent omic investigation of diatoms and Chlorophyta that secondarily lost photosynthesis, showed that those that retained a plastid genome have lost the genes encoding subunits of the complexes of the photosynthetic electron transport chain, but have maintained those encoding the subunits of the plastidial ATP synthase. It has been proposed that a functional plastid ATP synthase might be required in these algae to maintain a proton gradient across the thylakoid membranes via ATP hydrolysis. This pmf would be required by the twin arginine translocator (Tat) system for translocation of folded proteins into thylakoids (*57*). Indeed, the genes encoding the subunits of the Tat translocon being also conserved in these organisms. An inactivation of the Tat pathway in the Cyclotella *atpc* mutants would explain the conditional heterotrophic growth, by preventing the correct integration of Tat passengers such as the Rieske subunit of *b_6_f* complex, the PSBP and PSBQ subunits of the Oxygen Evolving Complex or the PSB27 and HCF136 PSII assembly factors…, but whether this is the case warrants further investigation.

To conclude, this study opens the way for detailed dissection of photosynthesis regulation and function in diatoms and proposes *C. cryptica* as a “brown *Chlamydomonas*” model system for secondary endosymbionts, with the potential to drive similarly transformative discoveries. As illustrated by the upregulated proteins identified in the γ-ATP synthase subunit knockout mutants, about 50% of proteins encoded by diatom genomes are still of unknown function, lacking orthologs outside of the bacilliarophyceae or stramenopile lineages. Inspired by the GreenCut inventory, originally based on *Chlamydomonas*(*78–80*), a catalog of nucleus-encoded proteins universally and specifically present in green algae and higher plants, but absent from non-photosynthetic species, *C. cryptica* could now be used to define a “BrownCut”: the set of nucleus-encoded, chloroplast-targeted proteins that are specific to diatoms (and possibly other secondary endosymbionts), but absent from archeplastidial red and green algae. By integrating genomic investigations, targeted mutagenesis, and biophysical analysis of photosynthetic phenotypes, *C. cryptica* offers a powerful platform to functionally characterize these diatom-specific proteins and advance our understanding of photosynthesis in the secondary endosymbiotic lineages that play a key role in aquatic carbon fixation and global ecosystems.

## METHODS

### Culture conditions

Axenic cultures of *C cryptica* CCMP332, obtained from the National Center for Marine Algae and Microbiota (NCMA Bigelow Laboratory for Ocean Sciences in East Boothbay, Maine, USA), were grown in artificial sea water medium (*81*) supplemented with L1 medium major nutrients and trace element solution (*82*). Cells were kept in vented culture flasks at 19°C in a constant light regime of 5, 25 or 50 µmol photons m^-2^ s^-1^. When stated, glucose (10 mM) was added to the medium in either light or dark cultures. Cell concentration was assessed using a Beckman Coulter Z1 S particle counter (Beckman Coulter, Brea, USA). For some experiments, photosynthesis was inhibited adding 1 µM DCMU (3-(3,4-dichloro-phenyl)-1,1-dimethylurea) directly into the culture medium.

### 13C-labelling experiments

Cells were first grown in L1 medium containing 10 mM of glucose in their respective conditions (5, 25, 50 µmol photons m^-2^ s^-1^ and dark) until reaching about 300.000 cell ml^-1^. Then they were washed three times in medium without glucose. Cells were diluted in medium containing 10 mM of 99% D-Glucose-2-^13^C (Cortecnet France, code CC855P1) to 30.000 cell ml^-1^ and incubated in flask of 100 ml with a working volume of 50 mL at the different light intensity conditions or in the dark. Each experiment was done in triplicate. Onday 4, cells were collected by centrifugation at 3000 g for 10 minutes at 4°C and washed three times in fresh medium. The pellets were frozen in liquid nitrogen and stored at −80°C until further analysis.

### Lipid extraction

Total lipid extraction was performed according to the Bligh and Dyer method (*83*) with some adjustments. Briefly, the cell pellet was extracted with a mixture of 3,8 ml of CHCl3/CH3OH/H2O 1:2:0.8 and sonicated for 5 minutes in ice bath. Supernatant was recovered by centrifugation at 3000 g for 5 minutes, 4°C. The extraction procedure was repeated twice. The organic phases were combined, and the solvent volumes were adjusted to obtain a mixture of CHCl3/CH3OH/H2O 2:2:0.8 allowing partitioning of organic and aqueous phase. The mixture was centrifuged at 1000 g for 5 min at 4 °C. The aqueous phase was discarded, and the underlying organic phase was evaporated to dryness under nitrogen. The dried lipid extract was transferred into a pre-weighed vial using a CHCl3/CH3OH (1:2) mixture, dried again and weighed to determine the total lipid content. The lipid extract was stored at −80°C until further analysis.

### Lipid analysis by LCMS

The lipid extracts were examined using a Vanquish ultra-high-performance liquid chromatography (UHPLC) system, equipped with a LUNA C18 column (5 μm particle size, 100 Å pore size, 150 mm × 2 mm; Phenomenex). The LC was connected to an Orbitrap Exploris 240 mass spectrometer (Thermo Scientific, Waltham, MA, USA), with a heated electrospray ionization (HESI) source. For each extract, 5 µL of a solution 50 µg/ml in CH3OH was injected. The chromatographic separation was conducted at a constant flow rate of 0.6 mL/min, with CH3OH and H2O (NH4OH/CH3COOH 0.005%, pH=8) as mobile phases A and B, respectively. The gradient elution began at 70% B, ramped linearly to 99% B over 30 minutes, held for column washing, and then returned to initial conditions for re-equilibration. HESI source was configured with the following parameters: The sheath gas flow was set to 40 arbitrary units (AU), the auxiliary gas at 5 AU, and the sweep gas flow at 2 L/min. The ion transfer tube and vaporizer temperatures were kept at 325°C and 300°C, respectively. The MS was operated in dual ionization mode with a spray voltage of 3.5 kV (positive mode) and −2.5 kV (negative mode). Mass spectra were acquired over an m/z range of 150–1500 with a resolving power of 60,000.

Quantitation of fatty acids was based on the relative abundance of MS fragments relative to m/z value of the entire molecules. Processing of raw data and peak integration was performed using Xcalibur and TraceFinder (Thermo). Relative quantitation of the different isotopologues of each fatty acid was adjusted for the natural background abundance of each isotopologue to make an accurate determination of artificial 13C label incorporation. For this purpose, we used FluxFix (http://fluxfix.science), an application freely available on the internet that automatically performs the calculation from raw signal intensity into percent mole enrichment for each isotopologue measured (*84*). To calculate total ^13^C enrichment of a molecule, percent mole enrichment was corrected for the number of ^13^C expected in each isotopologue.

### Genome editing of the *ATPC* gene

Mutagenesis of the *C. cryptica* nuclear-encoded *ATPC* gene (CCRYP_015031-RA in version 2 of the genome, g19148.t1 in the new annotation of the proteome by Nenasheva et al, 2025) was performed by CRISPR-Cas9 ribonucleoprotein (RNP) cell bombardment, by using *in vitro* pre-assembled RNP. Two gRNA sequences (gRNA1: 5’-GAATCCGTGCGTTACACCCC<u>TGG</u>-3’, gRNA2: 5’-CCGGAGAGGAC-GGCGAAACA<u>AGG</u>-3’; PAM sequence underlined) targeting the first exon and first intron of the *ATPC* gene (fig. Supp 6) were designed using Crispor (http://crispor.gi.ucsc.edu/) and verified with Cas-OFFinder (https://www.rgenome.net/cas-offinder/; (*85*). crRNAs and tracrRNA were purchased from Integrated DNA Technologies (IDT, Coralville, USA). The Alt-R^TM^ CRISPR-Cas9 System (IDT, Coralville, USA) was used following manufacturer’s instructions. Briefly, crRNA/tracrRNA duplex was assembled by mixing equal volumes of 100 µM solutions of each component on IDT duplex buffer and incubated at 95°C for 5 min and immediately cooled down at room temperature. RNPs were assembled by mixing crRNA/TracrRNA duplex with Alt-R^TM^ Cas9 enzyme, both diluted in phosphate buffer saline (PBS) to a final concentration of 6 µM and incubated at room temperature for 10-20 minutes. Two microliters of each RNP mix were precipitated onto 50 µl of 0.6 µm diameter gold particles (Bio-Rad, USA; 3 mg ml^-1^ suspension in water) together with 1 µg of the plasmid GK1074-pUC19-ccfcp6-nat, which contains the nourseothricin acetyl-transferase (*NAT*) gene flanked by the *C. cryptica* fucoxanthin chlorophyll a/c-binding protein 6 (*FCP*6) 5’- and 3’-UTRs. Furthermore, 2 µl of the cationic lipid polymer transfection reagent *Trans*IT^TM^ 2020 (Mirus bio, USA) were added to the gold particle RNP/plasmid mix and incubated for 10 minutes on ice as in (*86*) and then washed and resuspended in 50 µl of nuclease-free water. 10 ul of this suspension was used for each transformation by microparticle bombardment. Prior to bombardments, cells were grown in liquid ESAW medium to a cell density of 200 000 – 250 000 cells ml^-1^ and concentrated by centrifugation at 3 000g during 10 min. A total number of 1 x 10^8^ cells were plated in the center of 1.5% agar ESAW plates and let dry during 1 – 2 hours. Ten microliters of gold-RNP suspension were loaded in the center of a microcarrier disk and allowed to air dry. Biolistic bombardment was done using a PDS-1000/He^TM^ system (Bio-Rad, USA) with a rupture bursting pressure of 1 550 psi. After bombardment, cells were immediately scraped from the agar plate and transferred to 100 ml of non-selective liquid ESAW medium to let to recover in dim light for ∼16 hours without shaking. After that period, cells were plated at 5 x 10^6^ in agar ESAW plates containing 10mM glucose and 250 µg ml^-1^ nourseothricin (NTC) and kept in low light (≤5 µmol photons m^-2^ s^-1^) until colonies were observed (∼ 4 weeks).

Complemented lines were obtained by transforming a fragment of 2480 bp, amplified from the *C. cryptica* genomic DNA with the PCR primers Comp_FW and Comp_RV (Table S1), and containing the genomic *ATPC* gene and 691 bp upstream and 493 bp downstream of the start and stop codons, respectively. The fragment was transformed by microparticle bombardments as described above, and the complemented clones selected for their ability to rescue phototrophic growth on medium without glucose and under medium light (25 µmol photons m^-2^ s^-1^).

### Genotyping of mutant strains

Ten mL of exponentially growing wild-type, mutant and complemented lines were harvested by centrifugation (15 min at 3500 rpm, room temperature). Genomic DNA was extracted using the Invitrogen Easy-DNA gDNA Purification Kit, following Protocol #3 according to manufacturer’s instructions. The *ATPC* gene was amplified by PCR in a 25 µL reaction mixture containing: 1 µl DNA template, 1 µl dNTPs (10 µM), 1 µl primer mix (10 µM; see table S1), 0.2 µl GoTaq2 polymerase, 5 µL GoTaq2 buffer and nuclease free water to volume. PCR cycling conditions were: initial denaturation at 95°C for 2 min (preheated thermocycler), followed by 40 cycles of 95°C for 45 s, 55°C for 45 s and 72°C for 6 min, with a final extension at 72°C for 5 min. PCR products were purified and submitted to Eurofins Genomics for sequencing.

### Protein expression analyses

Protein expression analyses were performed on *C. cryptica* wild-type and three ATP synthase γ-subunit (*atpc*) mutant strains and a complemented light grown under continuous low light conditions (5 µmol photons m^-2^ s^-1^), with 10 mM glucose supplementation. For both western blot and proteomic analyses, 400 ml of cells in exponential phase of growth (around 3x 10^5^ cell.ml^-1^) were pelleted down at 8 000 rpm for 15 min. The pellets were washed in 40 mL PBS (phosphate buffered saline) solution and pelleted down at 8 000 rpm for 15 min. Pellets were resuspended in 300 µl of lysis buffer consisting of 30 mM Tris-Cl (pH 8), 50 mM NaCl, 1% SDS, 2 mM EDTA and cOmplete, EDTA-free, protease inhibitor cocktail from Roche (France). Cells lysis was performed by sonication on ice using a Vibra-Cell ultrasonic processor (Bioblock scientific) set to 40% amplitude. Samples were subjected to four cycles of 10 s pulses, each separated by 1 min intervals to prevent overheating. Lysis was confirmed by looking one lysate aliquot under the microscope. After lysis, cells were centrifuged at 13 500 rpm (max. speed) at 4°C during 30 min. The supernatant was recovered and protein concentration was measured with using the Pierce BCA protein assay kit (Thermo Fisher Scientific, USA) using bovine serum albumin (BSA) as protein standard.

Cell extracts were analyzed by SDS-PAGE on 8 to 16% or 12% acrylamide gels (Mini Protean Precast Gels, Bio-Rad) for immunoblot analysis after transfer to 0.1 µm nitrocellulose membranes (Amersham). Immunoblots were repeated at least twice. Proteins were detected by enhanced chemiluminescence using Clarity Western ECL Substrate (Bio-Rad). The dilutions used for the primary antibodies were 1:100,000 for anti-cytochrome *f*, anti-D1 and anti-PsaA, 1:80,000 for anti-RbcL, 1:50,000 for anti-CF1β and anti-CF1α, 1:20,000 for anti-D2, 1:10,000 for anti-AtpH and anti-ATPC and 1:8,000 for anti-PsaB. Detection was performed using horse-radish peroxidase (HRP)–conjugated secondary antibodies against rabbit IgG (Promega Corporation). Antibodies against the α and β subunits of F1/CF1 and cytochrome *f* have been de-scribed by Lemaire and Wollman (1989a)(*60*) and Kuras and Wollman (1994)(*87*), respectively. Antibodies against D1 (no. AS05 084), D2 (no. AS06 146), PsaA (no. AS06 172), RbcL (no. AS03 037), AtpH (no. AS09 591) and ATPC (no. AS08 312) were purchased from Agrisera.

### Mass spectrometry proteomic analysis

For proteomics, 20 µg of each protein sample were resuspended in Laemmli SDS buffer (Bio-Rad) and loaded on 12% polyacrylamide SDS-PAGE (Mini Protean Precast Gels, Bio-Rad). After a short 1.5 cm migration at 80V proteins were fixed and stained with Coomassie Brillant Blue R-250. Lanes containing proteins were excised manually and subjected to a manual in-gel digestion with modified porcine trypsin (Trypsin Gold, Promega). Briefly, after destaining, bands were subjected to a 30 minutes reduction step at 56°C in dark, 10 mM dithiotreitol in 50 mM ammonium bicarbonate (AMBIC) followed by a 1-hour cysteine alkylation step at room temperature in the dark with 50 mM iodoacetamide in 50 mM AMBIC. After dehydration under vacuum, bands were re-swollen with 250 ng of trypsin in 200 µL 50 mM AMBIC and proteins were digested overnight at 37 °C. Supernatants were kept and peptides present in gel pieces were extracted with 1% (v/v) trifluoroacetic acid and dried in a vacuum concentrator. Peptides were then solubilized in 100 µL of solvent A (0.1% (v/v) formic acid in 3% (v/v) acetonitrile) to a final concentration of 200 ng/µL. Tandem mass spectrometry of protein extracts were performed on a Q-Exactive Plus hybrid quadripole-orbitrap mass spectrometer (Thermo Fisher, San José, CA, USA) coupled to a Neo Vanquish liquid nano-chromatography (Thermoscientific). Peptide mixtures were analyzed in duplicate. Five microliters of peptide mixtures were loaded onto Neo pepmap trap column (300 µm x 5 mm, 5 µm, 100 Å; ThermoFisher Scientific) equilibrated in solvent A and separated at a constant flow rate of 300 nl/min on a PepMap NEO™ RSLC C18 Easy-Spray column (75 µm x 50 cm, 2 µm, 100 Å; Thermo Scientific) with a 90 min gradient (0 to 20% B solvent (80%ACN, 0.1% formic acid (v/v)) in 60 min and 20 to 35% B solvent in 30 min).

Data acquisition was performed in positive and data-dependent modes. Full scan MS spectra (mass range m/z 400-1600) were acquired in profile mode with a resolution of 70,000 (at m/z 200) and MS/MS spectra were acquired in centroid mode at a resolution of 17,500 (at m/z 200). All other parameters were kept as described in (*88*).

### Data processing, quantification and differential expression analysis

Raw data were processed using the MaxQuant software package (http://www.maxquant.org, version 1.5.6.5) (*89*). Protein identifications and target decoy searches were performed using the Andromeda search engine and an home-made database (31572 entries) containing all genes models from the new annotation of the proteome by Nenasheva et al, 2025 (*36*), plus all organelle-encoded proteins and the selection marker (NAT) in combination with the Maxquant contaminants database (number of contaminants: 245). The mass tolerance in MS and MS/MS was set to 10 ppm and 20 mDa, respectively. Methionine oxidation and protein N-term acetylation were taken into consideration as variable modifications whereas cysteine carbamidomethylation was considered as fixed modification. Trypsin was selected as cutting enzyme and 2 missed cleavages were allowed. Proteins were validated if at least 2 unique peptides having a protein FDR < 0.01 were identified. The setting “Match between runs” was also taken into consideration to increase the number of identified peptides. For quantification, we used unique and razor peptides with a minimum ratio count ≥ 2 unique peptides. Protein intensities were calculated by Delayed Normalization and Maximal Peptide Ratio Extraction (MaxLFQ) (*90*).

For differential expression analysis, the proteinGroup.txt output from MaxQuant was imported into R version 4.5.1 (*91*)) for downstream processing and statistical analysis. Contaminant proteins, reverse sequences and proteins identified “only by site” were removed. Proteins were further filtered to retain only those with at least two valid values in at least one condition. Label-free quantification (LFQ) values were then log2-transformed and missing values were imputed under a ‘Missing Not At Random’ (MNAR) assumption by sampling from a left-shifted Gaussian distribution (1.8 standard deviations, width 0.3). Differential expression analysis was carried out by comparing each mutant strain to the wild-type using protein-wise linear modelling with empirical Bayes moderation. Differentially expressed proteins were identified with the *limma* package v3.64.1 (*92*). Proteins were considered deregulated if they met two criteria: an adjusted p-value (Benjamini-Hochberg correction) < 0.05 and an absolute log2 fold change ≥ 1. Finally, the subcellular localization of all identified proteins was predicted with ASAFind (*93*).

### Electron microscopy

Cells were fixed in 2% glutaraldehyde in Filtered Sea Water (FSW) for 2 hours on ice and then washed three times in FSW and three times in ddH2O. Between each washing step of 15 minutes cells were centrifuged at 200g for 10 minutes at 4°C. Cells were then post-fixed in 1% osmium tetroxide (OsO4) and 1.5% Potassium ferrocyanide for 1hour on ice, followed by an incubation in 0.3% Thiocarbohydrazide at room temperature for 20 minutes and then in 1% OsO4 for 1 hour at RT. Cells were washed three times in between each post-fixation step in ddH2O for 15 minutes. After the last washing step, the pellets were embedded in 0.7% ultra-low gelling temperature agarose and dehydrated in a graded ethanol series further substituted by propylene oxide and embedded in Epon 812 at room temperature and polymerized at 60°C for 3 days. Resin blocks were sectioned with a Ultracut UCT ultramicrotome (Leica, Vienna, Austria), ultrathin sections (70 nm) were placed on nickel grids and observed with a Zeiss LEO 912AB EFTEM (Zeiss, Oberkochen, Germany).

### Chlorophyll fluorescence and spectroscopic measurements

*C. cryptica* cultures were sampled during the exponential growth phase, unless otherwise indicated (e.g., for samples measured after several days in the dark). Prior to measurements, samples were concentrated approximately 10-fold by centrifugation (3 000 × g, 10 min), then allowed to recover for 5 to 10 minutes under gentle shaking and very low light (∼2 μmol photons m⁻² s⁻¹). Fluorescence parameters were measured using a SpeedZen fluorescence imaging system (JbeamBio, France), which allows the simultaneous analysis of multiple 60 μL samples. The system uses green actinic light and saturating pulses at 532 nm. For dark-adapted samples, minimum fluorescence (F₀) and maximum fluorescence (Fₘ) were measured before and after a 250 ms saturating pulse of 5,000 μmol photons m⁻² s⁻¹, respectively. These values were used to calculate the maximum quantum yield of PSII, as follows: Fᵥ/Fₘ = (Fₘ - F₀)/Fₘ To assess the dependence of photosynthesis on light intensity, measurements were performed under increasing actinic light intensities ranging from 35 to 600 μmol photons m⁻² s⁻¹. Each light intensity was applied for 5 minutes, with a saturating pulse every minute. Under stable light conditions, steady-state fluorescence (Fₛ) and maximum fluorescence in the light (Fₘ′) were measured before and after each saturating pulse and used to calculate: PSII operating efficiency, <λ_PSII_ = (Fₘ′ − Fₛ) / Fₘ′ and Non-photo-chemical quenching (NPQ), NPQ = (Fₘ − Fₘ′) / Fₘ′. Relative electron transport rate (rETR) was calculated as rETR = <λ_PSII_ × light intensity

For transient absorption measurements, experiments were conducted in a JTS-10 spectrophotometer (Biologic, France), allowing to measure absorption change signals at different wavelengths by placing specific interference filters (3-8 nm bandwidth) on the detecting light path. Several absorption change signals were probed: the electrochromic shift (ECS) of pigments absorption spectrum and absorption changes associated to oxidized-minus-reduced signals from the electron carriers between cytochrome b6f and PSI, namely P700 and c-type cytochromes. The c-type cytochromes (cyt.) and ECSlin signals were obtained through measuring transient absorption changes at 520, 554 and 573 nm with BG39 filter on the measuring and reference photodiodes (Schott (Mainz, Germany)) as in [8]. Because of slight differences in the spectra of linear and quadratic ECS, signals of interest were then calculated as cyt. = [554]-[520] in the WT, or [554]-1.25*[520]+0.2*[573] in *atpc* mutants. Then linear ECS was calculated as ECSlin = [520]-0.25*cyt. in all strains. Absorption difference signals following a saturating single turnover flash from a dye laser (690 nm) were used as internal normalization: they provided the ECSlin signal corresponding to one single charge separation (CS) per PS, and the oxidation of 1 cyt. per PSI. PSI/PSII stoichiometry was measured through the comparison of the ECSlin signal induced by a saturating laser flash without (PSII + PSI) or with 10 µM 3-(3,4-dichlorophenyl) 1,1–dimethylurea (DCMU) and 300 µM hydroxylamine (HA) (PSI alone). Absorption difference signals related to the redox changes of P700 were measured at 705 nm – 730 nm with a high pass RG695 filters on the measuring and reference photodiodes (the difference between the two wavelengths allows to eliminate for the contribution of ferredoxin and the scattering signals). The maximal absorption change signal corresponding to full oxidation of P700 (PM) was then measured at the end of a 20 ms saturating pulse (red light, 660 nm, 6000 µmol photons m^-2^ s^-1^) in the presence of 10 µM 3-(3,4-dichlorophenyl) 1,1–dimethylurea (DCMU). The light dependence of PSI yield was recorded at actinic light intensities ranging from 3 to 350 µmol photons m^-2^ s^-1^. PSI yield was calculated as <λ_PSI_ = (Pm’ – P)/Pm, where P is the steady-state absorption under actinic light and Pm’ is the maximal absorption after a saturating light pulse under actinic light. The proportion of oxidized P_700_ was calculated as (P – P0)/(Pm– P0), were P0 is the absorption in the dark-adapted sample. Furthermore, the effect of luminal pH on P700 oxidation was determined by measuring <λ_PSI_ at 56 µmol photons m^-2^ s^-1^ in the presence of 1 µM nigericin.

## Acknowledgements

This work was supported by ANR BrownCut (ANR-19-CE20-0020) and the European Union’s Horizon Europe Programme BlueRemediomics (grant agreement no. 101082304; views and opinions expressed are those of the author(s) alone and do not necessarily reflect those of the European Union or the European Research Executive Agency. Neither the European Union nor the granting authority can be held responsible for them) to A.F, the LABEX DYNAMO (ANR-LABX-011), EQUIPEX (CACSICE ANR-11-EQPX-0008) and CPER-équipement PSL-RESOLUTION notably through funding of the Proteomic Platform of IBPC (PPI). This project also received funding from the European Union’s Horizon 2020 research and innovation programme under the Marie Skłodowska-Curie grant agreement No 101034407 COFUND FP-YNAMO-PARIS). We thank Dr. Nicole Poulsen for kindly providing the transformation vector GK1074-pUC19-ccfcp6-nat used in this work.

---

**Fig. S1.**
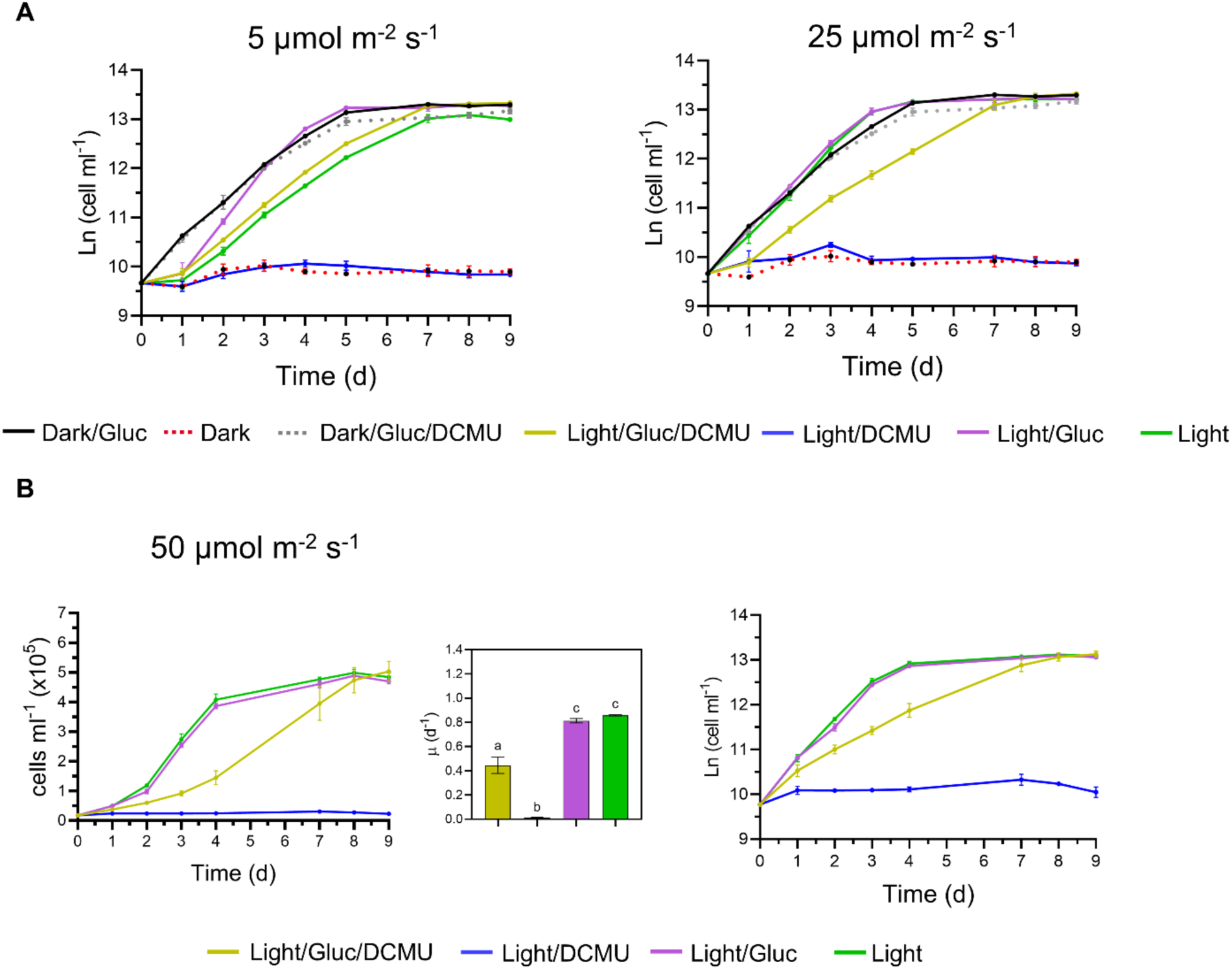
(**A**) growth curves from Fig. 1A shown in logarithmic scale. (**B**) growth curves in linear (left) and logarithmic (right) scale of *C. cryptica* wild type grown at 50 µmol photons m^-2^ s^-1^. Graphs are shown as the mean ± SD of three biological replicates. For statistics one-way ANOVA was used, with Tukey’s post-hoc analysis for multiple comparisons (different letters represent significant differences, p≥0.05).

**Fig. S2.**
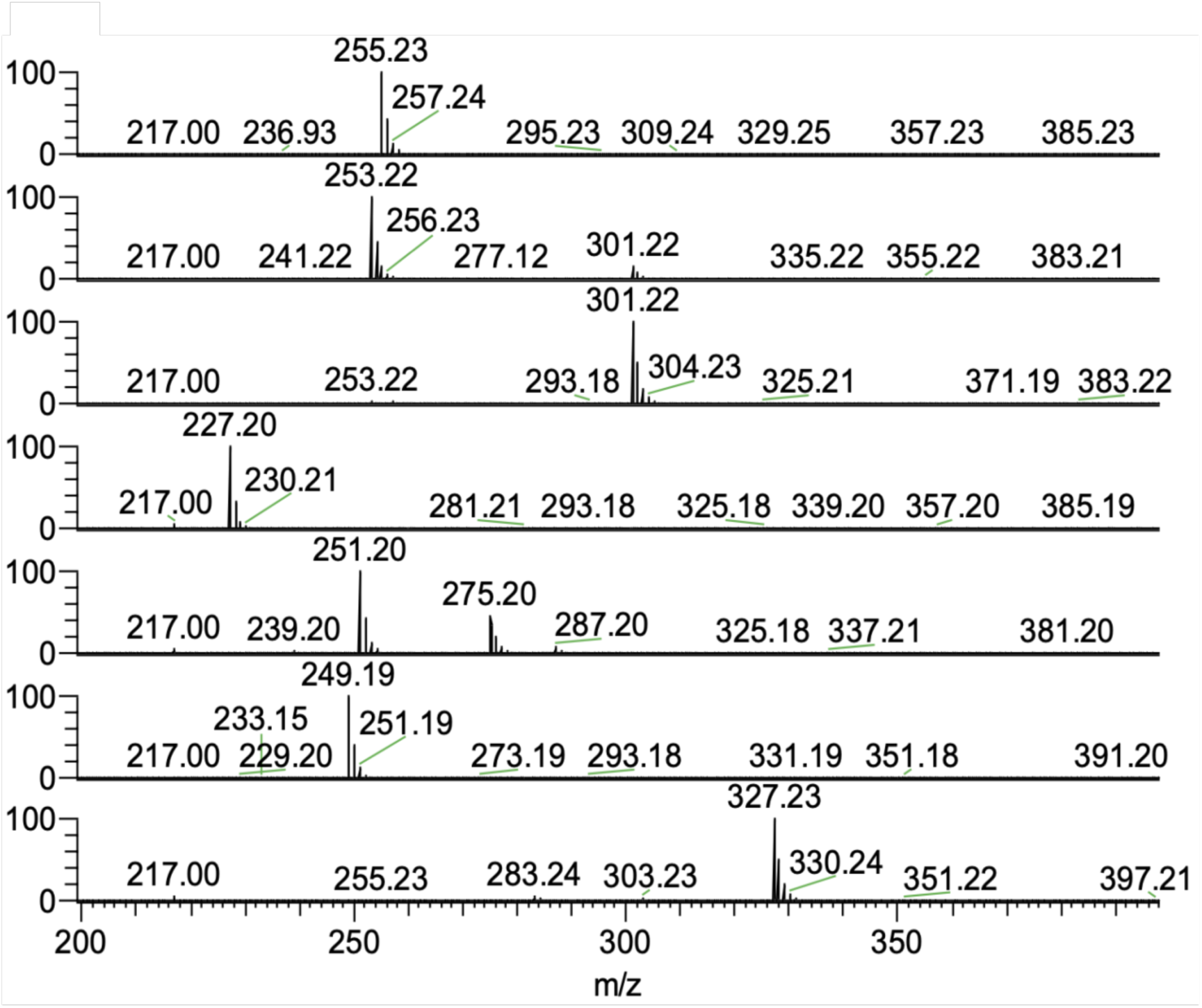
^13^C labelling of major fatty acids (above 10% threshold) of *C. cryptica* grown at 50 µmol photons m^-2^s^-1^ and 2-^13^C-glucose.

**Fig. S3.**
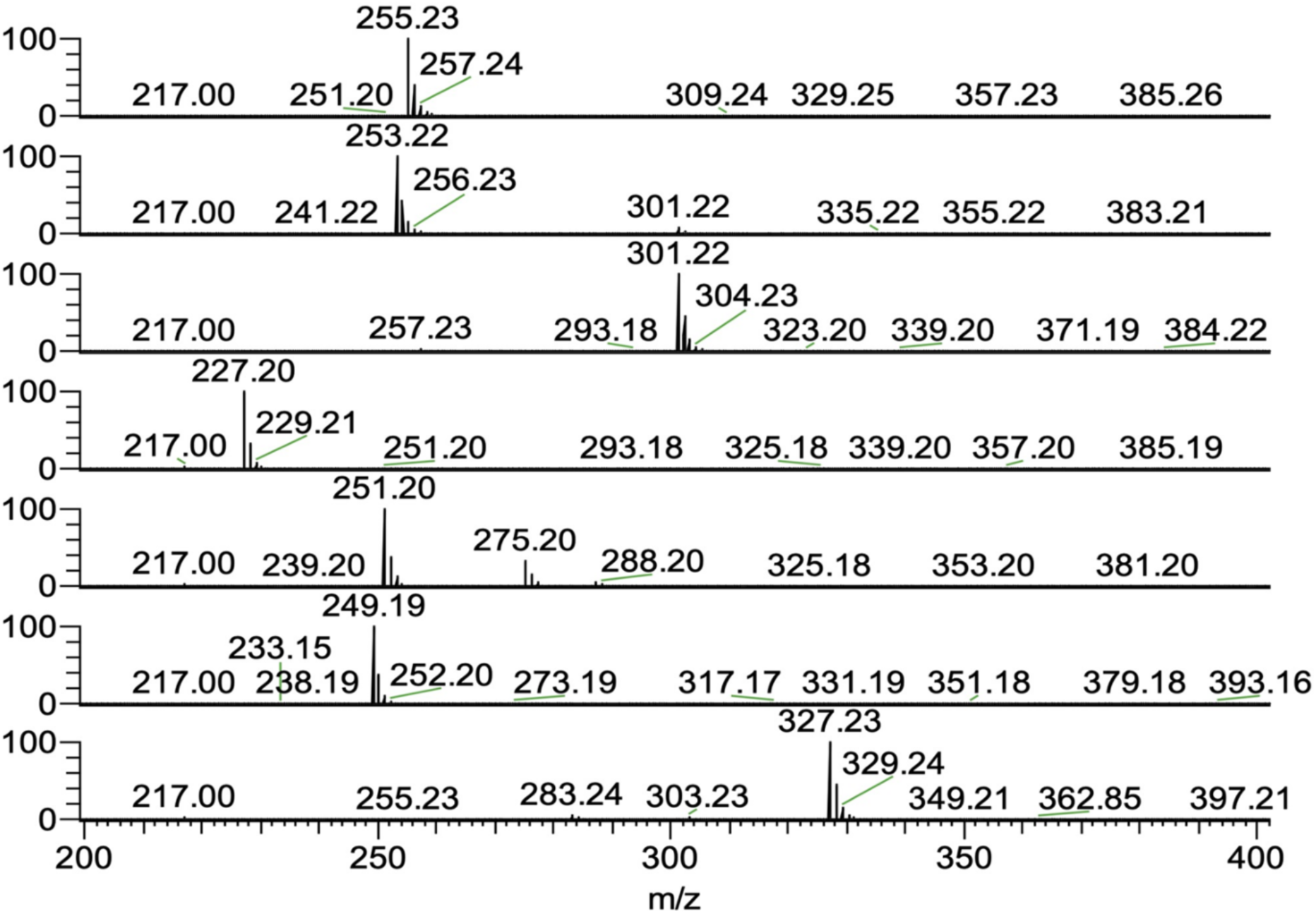
^13^C labelling of major fatty acids (above 10% threshold) of *C. cryptica* grown at 25 µmol photons m^-2^s^-1^ and 2-^13^C-glucose.

**Fig. S4.**
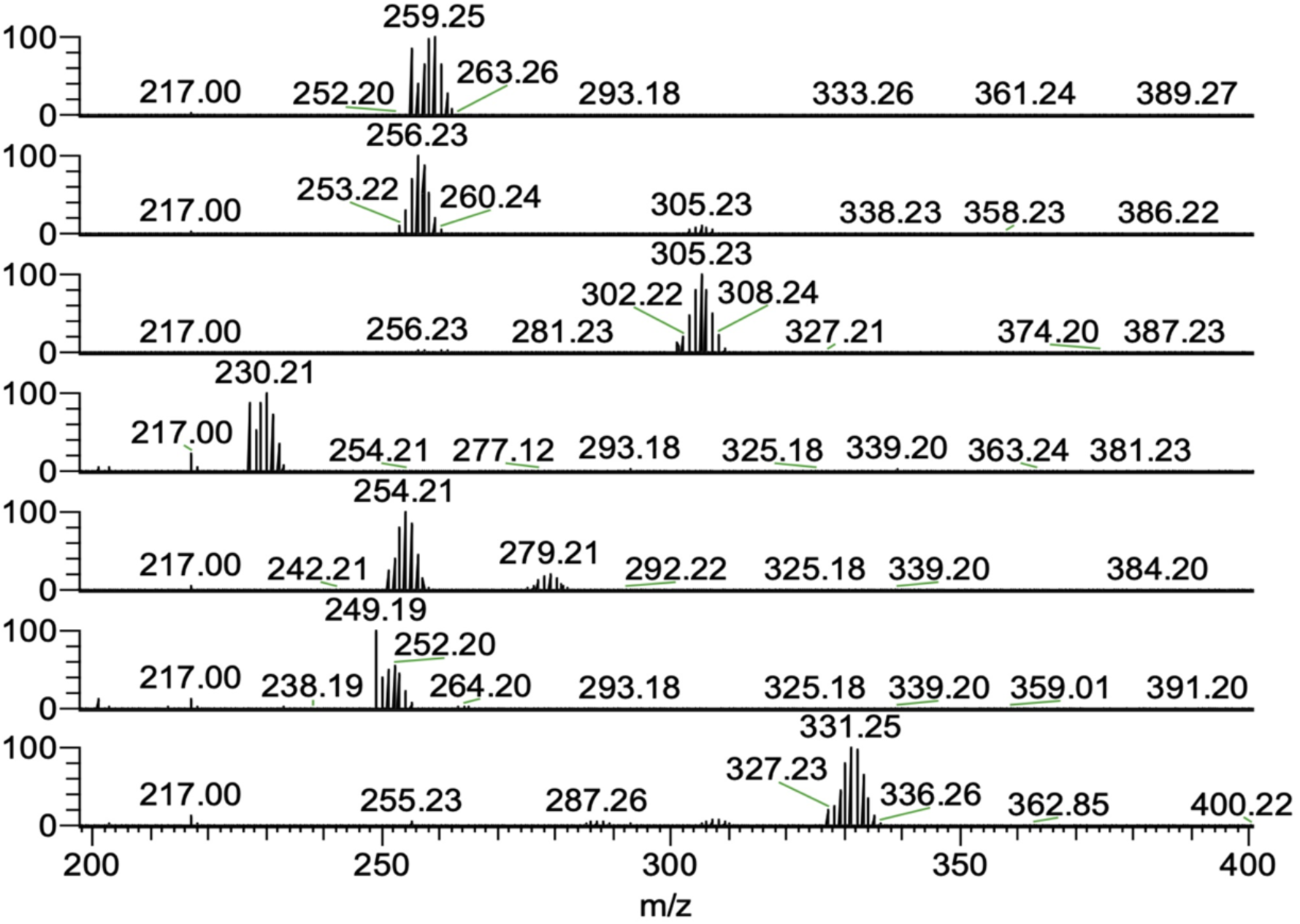
^13^C labelling of major fatty acids (above 10% threshold) of *C. cryptica* grown at 5 µmol photons m^-2^s^-1^ and 2-^13^C-glucose.

**Fig. S5.**
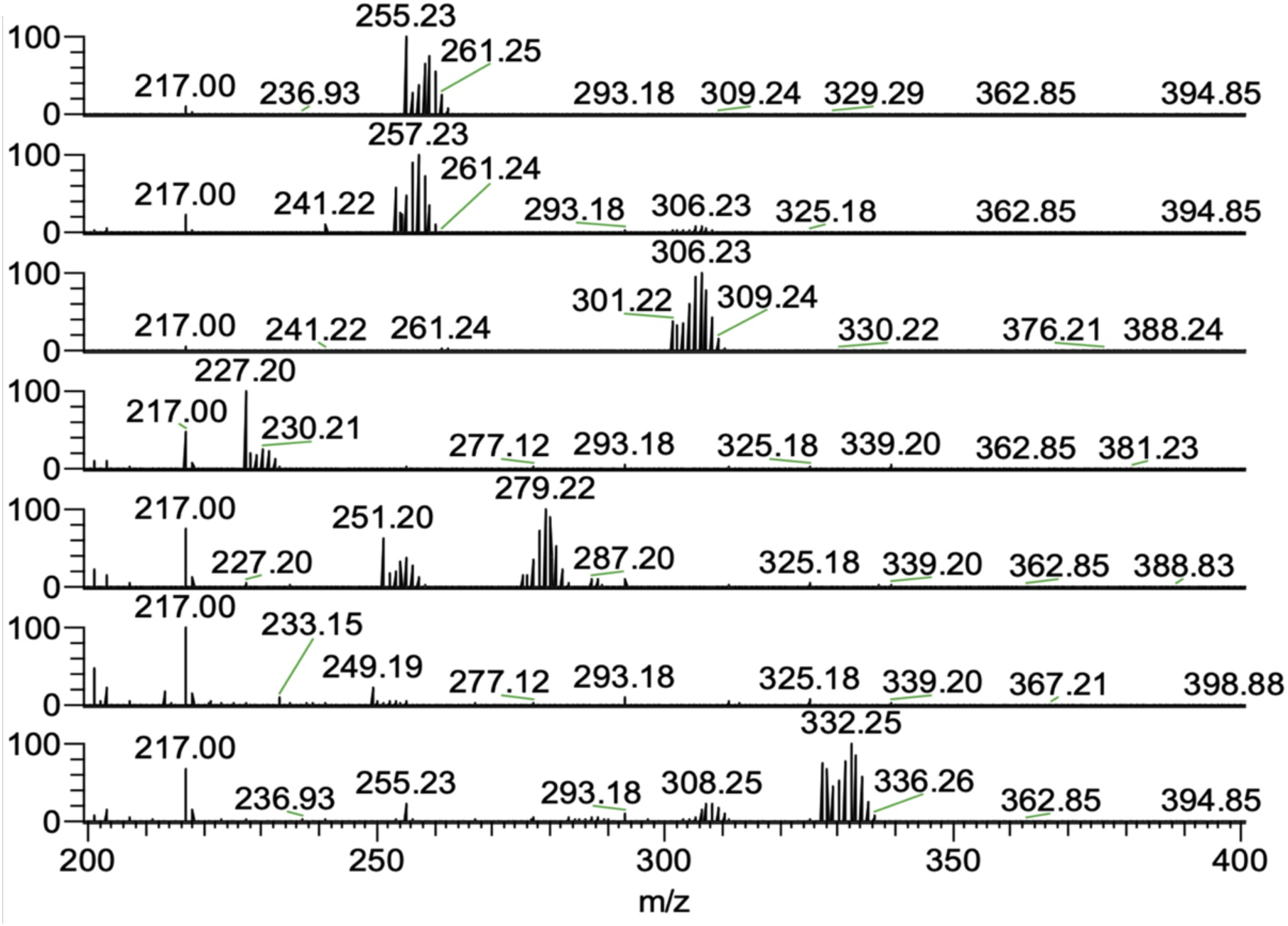
^13^C labelling of major fatty acids (above 10% threshold) of *C. cryptica* grown in the precense of 2-^13^C-glucose in complete darkness.

**Fig. S6.**
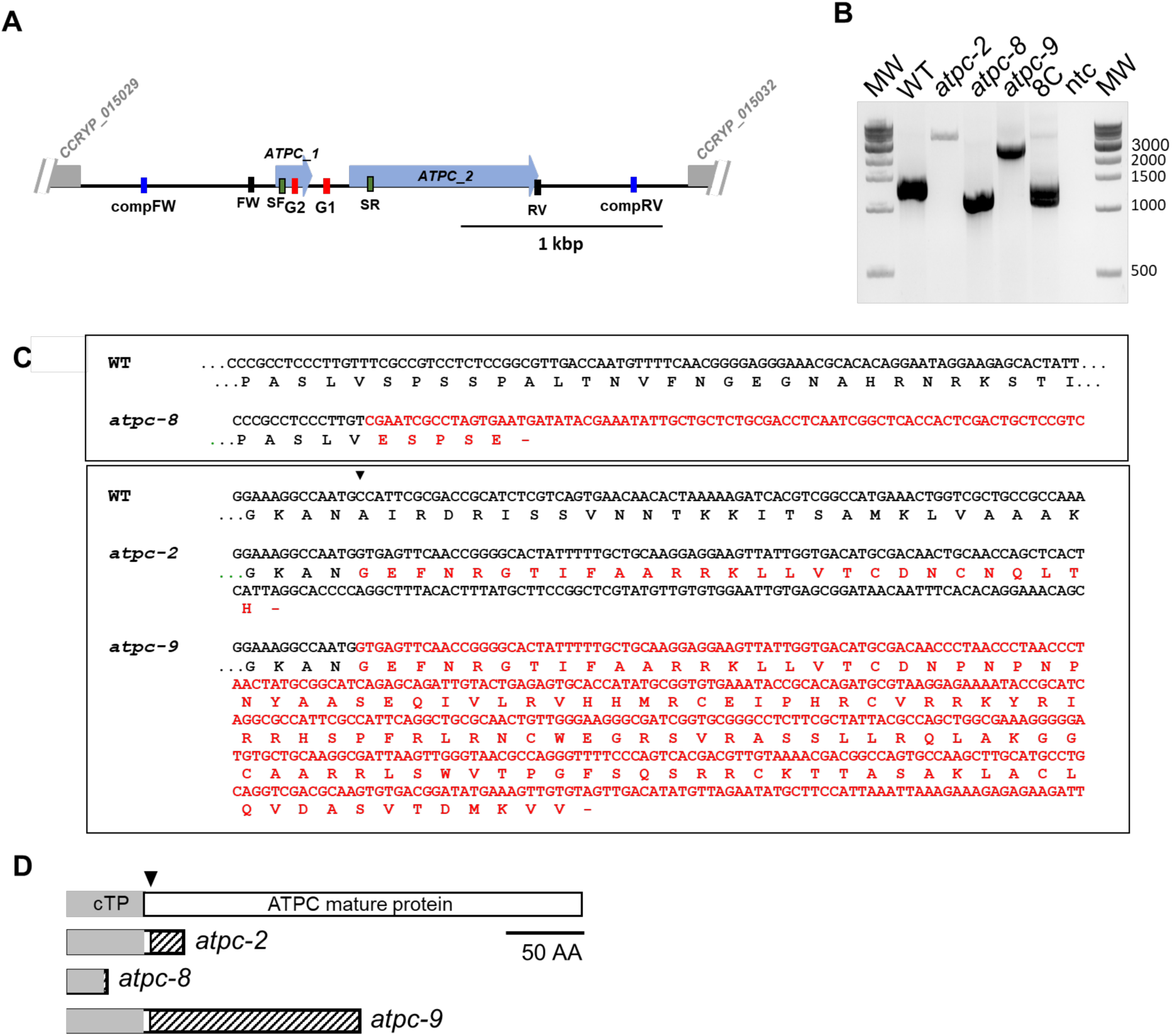
Mutations at the *ATPC* locus. **(A)** schematic representation of the *ATPC* locus: exons are shown as blue arrows while the beginning of the first exon of the adjacent genes is depicted as grey boxes. The position of the oligonucleotides used to amplify the region surrounding the expected deletion in panel B (FW and RV, black lines), the fragment used for the complementation (compFW and compRV, blue lines) and the primer pair used for screening of complemented strains (SF and SR) is shown as well as that of the two guide RNAs (G2 and G1 red lines). (**B**) PCR amplification of the *ATPC* gene in the wild type, the three *atpc* mutant strains and in the complemented strain 8C; MW molecular weight marker, ntc: no template control. (**C**) Comparison of the wild-type *ATPC* sequence with those in the *atpc* mutant and consequences of the mutations on translation. Nucleotides and amino acids that differ in the mutant from those found in the wild type are written in red. Upper: WT vs. mutant strain *atpc*-8; Lower: WT vs *atpc*-2 and *atpc*-9 mutant strains. The arrowhead points to the junction boundary in the Wild-type. (**D**) Schematic representation of what is shown panel C.

**Fig. S7.**
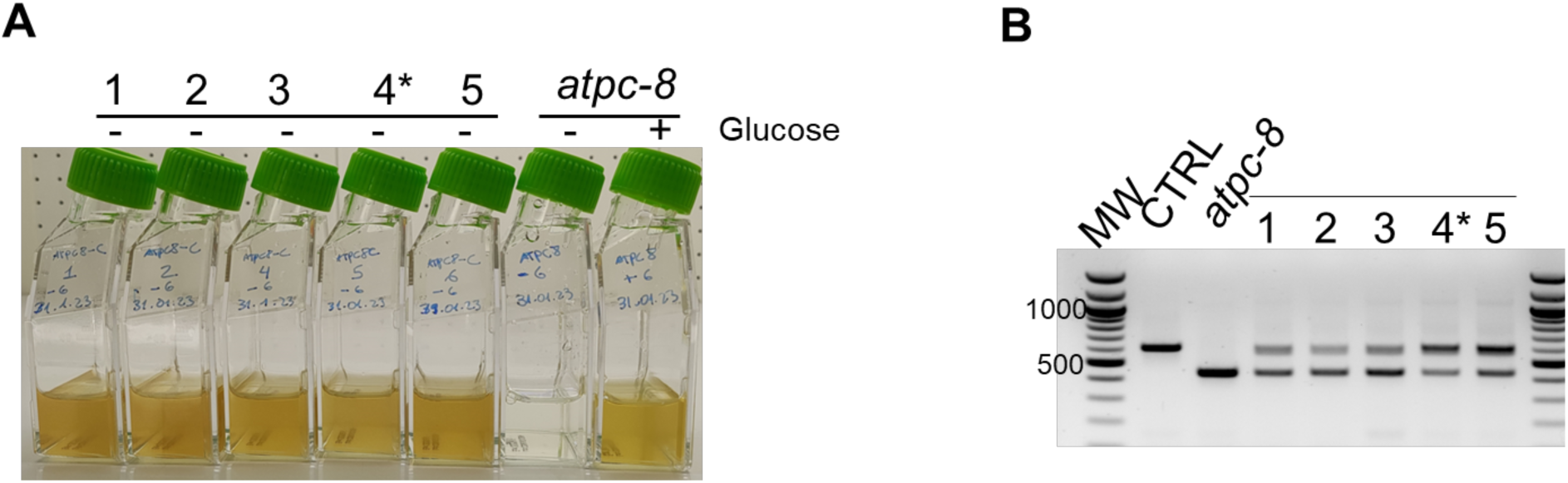
(**A**) Growth of the complemented strains (1–5) in the absence of glucose showing the recovery of phototrophy. The recipient strain *atpc-*8 is shown for comparison. Asterisk denotes the complemented strain selected to be used in subsequent experiment, named 8C. (**B**) PCR amplification of the *ATPC* locus in the complemented strains, in the wild type and the *atpc-*8 recipient strain for comparison using primers SF and SR (Suppl. Fig. 2). Complementation was confirmed by the presence of the WT PCR product (599 bp compared to 428 bp in the *atpc*-8) in all the five strains. Both the wild-type and the mutant amplicons are observed.

**Fig. S8.**
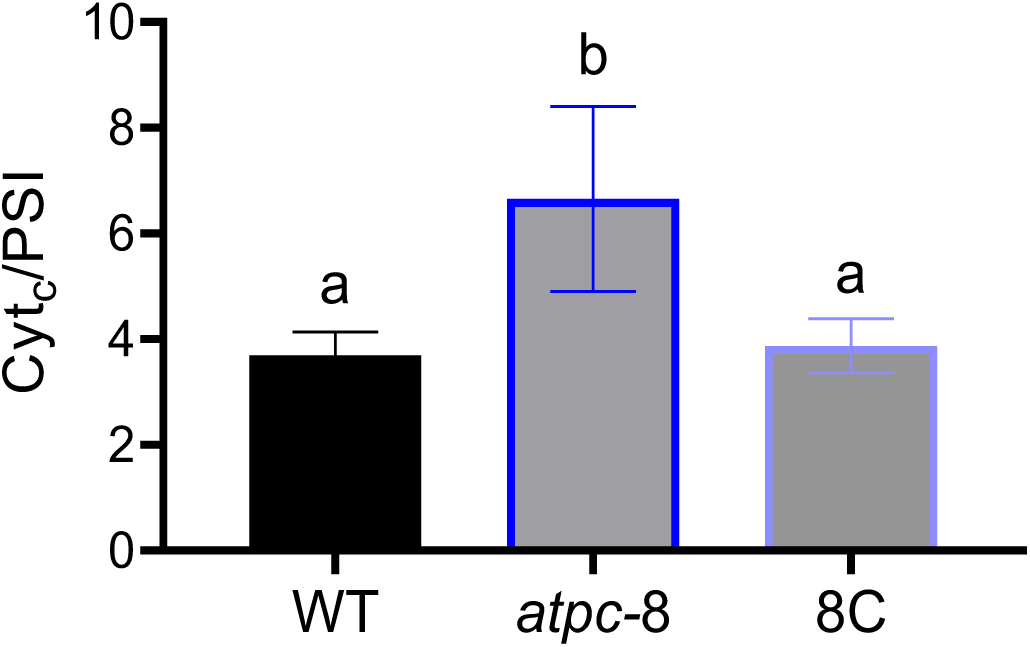
c-Type cytochromes oxidation to PSI obtained from ECS data. Graph show the average ± SD of three biological replicates. For statistics, one-way ANOVA was used, with Tukey’s post-hoc analysis for multiple comparisons where different letters represent significant differences, p≥0.05.

**Fig. S9.**
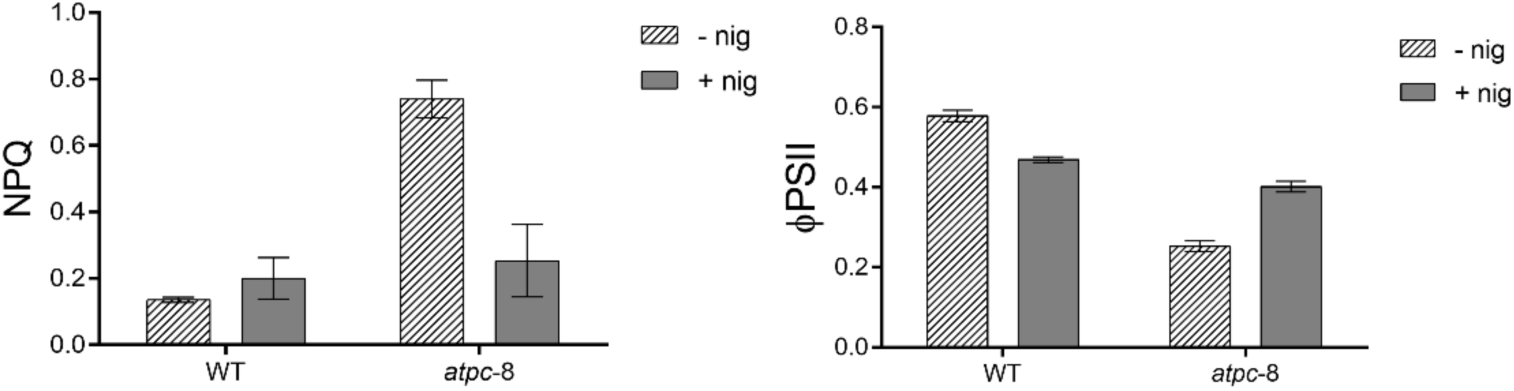
effect of 1 µM nigericin on NPQ (left) and ΦPSII (right) in WT and *atpc-*8 mutant. As consequence of the dissipation of ΔpH, NPQ is relaxed in the *atpc*-8 mutant and electron flow in PSII increased. Measurements were made at a light intensity of 35 µmol photons m^-2^ s^-1^. Graphs are shown as the mean ± SD of three biological replicates

**Table S1.**
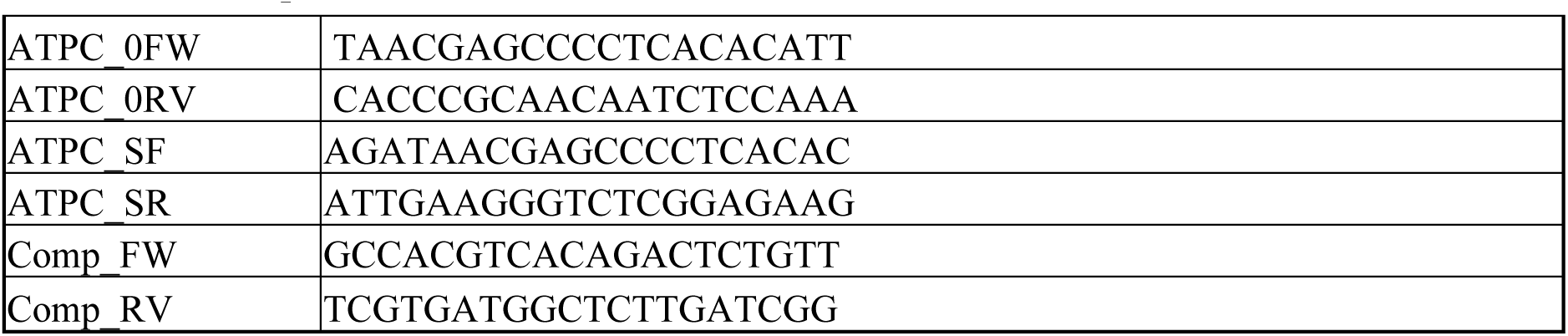
List of primers used in this work.

**Table S2.** ***C. cryptica* culture medium composition.**

| **Component** | **Final Concentration (M)** | **Final Concentration (g/L)** |
| --- | --- | --- |
| **Salt Solution 1 (1X)** |  |  |
| NaCl | 0.363 M | 21.2 g/L |
| Na<sub>2</sub>SO<sub>4</sub> | 25.0 mM | 3.55 g/L |
| KCl | 8.0 mM | 0.599 g/L |
| NaHCO<sub>3</sub> | 2.07 mM | 0.174 g/L |
| KBr | 0.725 mM | 0.086 g/L |
| H<sub>3</sub>BO<sub>3</sub> | 0.372 mM | 0.023 g/L |
| NaF | 67 µM | 0.0028 g/L |
| **Salt Solution 2 (1X)** |  |  |
| MgCl<sub>2</sub>·6H<sub>2</sub>O | 47.2 mM | 9.60 g/L |
| CaCl<sub>2</sub>·2H<sub>2</sub>O | 9.15 mM | 1.34 g/L |
| SrCl<sub>2</sub>·6H<sub>2</sub>O | 82 µM | 0.0218 g/L |
| **Macronutrients** |  |  |
| NaNO<sub>3</sub> | 0.882 mM | 0.075 g/L |
| NaH<sub>2</sub>PO<sub>4</sub>·H<sub>2</sub>O | 36 µM | 0.005 g/L |
| Na<sub>2</sub>SiO<sub>3</sub>·9H<sub>2</sub>O | 0.21 mM | 0.060 g/L |
| FeEDTA | 6.8 µM | 0.0025 g/L |
| **Trace Metals** |  |  |
| Na<sub>2</sub>EDTA·2H<sub>2</sub>O | 8.3 µM | — |
| ZnSO<sub>4</sub>·7H<sub>2</sub>O | 0.25 µM | — |
| CoCl<sub>2</sub>·6H<sub>2</sub>O | 67 nM | — |
| MnCl<sub>2</sub>·4H<sub>2</sub>O | 2.7 µM | — |
| Na<sub>2</sub>MoO<sub>4</sub>·2H<sub>2</sub>O | 6 nM | — |
| Na<sub>2</sub>SeO<sub>3</sub> | 1 nM | — |
| NiCl<sub>2</sub>·6H<sub>2</sub>O | 6 nM | — |
| CuSO<sub>4</sub>·5H<sub>2</sub>O | 39 nM | — |
| **Vitamins** |  |  |
| Thiamine (B1) | 11.9 nM | — |
| Vitamin B12 | 14.8 pM | — |
| Biotin | 8.2 pM | — |

